# The E3 ligase HECTD4 regulates COX-2 dependent tumor progression and metastasis

**DOI:** 10.1101/2025.04.09.647907

**Authors:** Joanna A. Vuille, Cem Tanriover, Douglas S. Micalizzi, Richard Y. Ebright, Sambhavi Animesh, Robert Morris, Soroush Hajizadeh, Zachary J. Nicholson, Hunter C. Russell, Eric F. Zaniewski, Ben S. Wittner, Ben K. Wesley, Julian Grünewald, Regan N. Szalay, Ezgi Antmen, Douglas B. Fox, Min Yang, J. Keith Joung, Doga C. Gulhan, Andrew E.H. Elia, Wilhelm Haas, Eugene Oh, Shyamala Maheswaran, Daniel A. Haber

## Abstract

E3 ubiquitin ligases mediating turnover of proteins engaged in cancer progression point to key regulatory nodes. To uncover modifiers of metastatic competency, we conducted an in vivo genome-wide CRISPR-inactivation screen using cultured breast circulating tumor cells, following intravascular seeding and lung colonization. We identified HECTD4, a previously uncharacterized gene encoding a conserved potential HECT domain-containing ubiquitin transferase, as a potent tumor and metastasis suppressor. We show that purified HECTD4 mediates ubiquitin conjugation in vitro, and proteomic studies combined with ubiquitin remnant profiling identify a major degradation target as the prostaglandin synthetic enzyme cyclooxygenase-2 (COX-2; PTGS2). In addition to COX-2 itself, HECTD4 targets its regulatory kinase MKK7. In breast cancer models, HECTD4 expression is induced as cells lose adherence to the matrix, and its depletion massively increases COX-2 expression, enhancing anchorage-independent proliferation and tumorigenesis. Genetic or pharmacologic suppression of COX-2 reverses the pro-tumorigenic and pro-metastatic phenotype of HECTD4-depleted cells. Thus, HECTD4 encodes an E3 ubiquitin ligase that downregulates COX-2 suppressing anchorage-independence in epithelial cancer cells.

**Significance Statement:** A genome-wide CRISPR-inactivation screen identified the previously uncharacterized E3 ubiquitin ligase HECTD4, as a tumor and metastasis suppressor, with COX-2 as its major degradation target. The pro-tumorigenic and pro-metastatic effect of HECTD4 suppression depends on COX-2 stabilization, which is critical for anchorage-independent growth, providing a basis for investigating COX-2 inhibition to prevent metastatic recurrence.

## Introduction

Circulating tumor cells (CTCs) are shed from cancerous lesions into the bloodstream and constitute precursors of metastasis. These circulating cancer cells exhibit the heterogeneity acquired by advanced tumors, as they initially respond and then progress following therapeutic interventions by adapting to loss of matrix adhesion, high oxygen tensions and circulatory stress in the bloodstream (1). By using a microfluidic platform to deplete leukocytes from blood samples and preserve viable CTCs, we have successfully established long-term cultures of breast cancer CTCs, maintained under anchorage-independent, hypoxic, stem cell culture conditions (2, 3). While cultured CTCs produce orthotopic mammary gland tumors in immunosuppressed mice, they remarkably fail to initiate lung tumors following direct tail vein intravenous inoculation (4). Identifying factors that modulate the functional properties of cultured CTCs may, thus, identify strategies to suppress blood-borne cancer metastasis.

Inflammatory mediators secreted into the microenvironment may mediate both autocrine and microenvironmental effects on tumorigenesis. The cyclooxygenase COX-2, itself triggered by inflammatory stimuli, catalyzes the synthesis of prostaglandins such as PGE2, contributing to cancer cell proliferation, apoptosis resistance, angiogenesis and fibrosis (5, 6). Consistent with an important role in tumor initiation, seminal epidemiological studies of the COX-2 inhibitor Celecoxib demonstrated striking benefits in colorectal cancer prevention, but at the risk of increased thrombosis and cardiovascular events, leading to the discontinuation of cancer prevention studies (7, 8). A role for COX-2 suppression of metastasis through non-steroidal anti-inflammatory drugs (NSAID) has also been suggested in epidemiological studies (9), but it is unclear whether this reflects tumor cell-specific effects, anti-inflammatory activity or alterations in platelet aggregation. Recent and ongoing clinical studies have focused on aspirin use to suppress metastatic recurrence in patients with a history of cancer, with ongoing consideration of predictive biomarkers, patient stratification, drug dosage and duration of follow up (9–11). The potential utility of COX-2 inhibitors in the suppression of metastasis thus remains to be resolved.

Here, we conducted a genome-wide *in vivo* CRISPR-inactivation (CRISPR-i) screen using cultured breast CTCs, identifying *HECTD4,* a previously uncharacterized member of the HECT ubiquitin ligase gene family, as a suppressor of tumor initiation in a model of lung metastasis. We show that HECTD4 specifically promotes the ubiquitination and degradation of COX-2, an effect that is most dramatic as breast cancer cells lose cell adhesion. *COX-2* knockdown within breast cancer cells abrogates the anchorage-independent survival benefit of *HECTD4* suppression, consistent with a tumor cell-intrinsic role for COX-2 activity in metastasis initiation.

## Results

### Suppression of CTC-mediated metastasis by HECTD4 in a CRISPR-i screen

To identify negative regulators of CTC-mediated metastasis, we introduced a genome wide CRISPR-i library into BRx-142 cells, a CTC-derived cell line generated from a blood sample of a patient with advanced, refractory hormone receptor-positive (HR+) breast cancer (3). BRx-142 CTCs are highly tumorigenic following orthotopic inoculation in NSG immunosuppressed mice, yet they fail to produce sustained metastases in the lungs following intravenous tail vein inoculation (4). The cultured CTCs were first transduced to stably express a catalytically inactive Cas9 (dCas9) fused to a transcription repressor domain (KRAB), followed by transduction with a sgRNA library targeting > 18,000 genes, with 3 guides per gene at a MOI of 0.3, to achieve a representation of 300-350 cells per sgRNA (12). NSG immunodeficient mice (n=15) were inoculated by tail vein with the engineered Brx142 CTCs and their lungs were harvested after 3 months, followed by bulk sequencing to score sgRNA enrichment, compared with sample input (**FIG. 1A**). To mitigate the heterogeneity of guide representation, we pooled sequences from all mice, excluded low-input sgRNAs with fewer than 100 absolute reads in the viral pool (148 sgRNAs) and rank-ordered genes based on their most highly enriched gRNA (see Methods). Among the high-scoring genes were known mediators of tumor progression, including genes involved in transcription/translation (n = 35) and apoptosis (n = 21) (**FIG. 1B**). Remarkably, multiple hits identified genes implicated in proteolytic degradation, with seven E3 ubiquitin ligases ranking among the top 250 hits. Six of these genes encode the E3 ligases *FBXL6, WWP1, UHRF2, KCMF1, JADE2 and RNF121,* each with previously well-defined substrates and a seventh (ranked 49), *HECTD4*, whose substrate and function is unknown (**FIG. 1C**).

**Figure 1.**
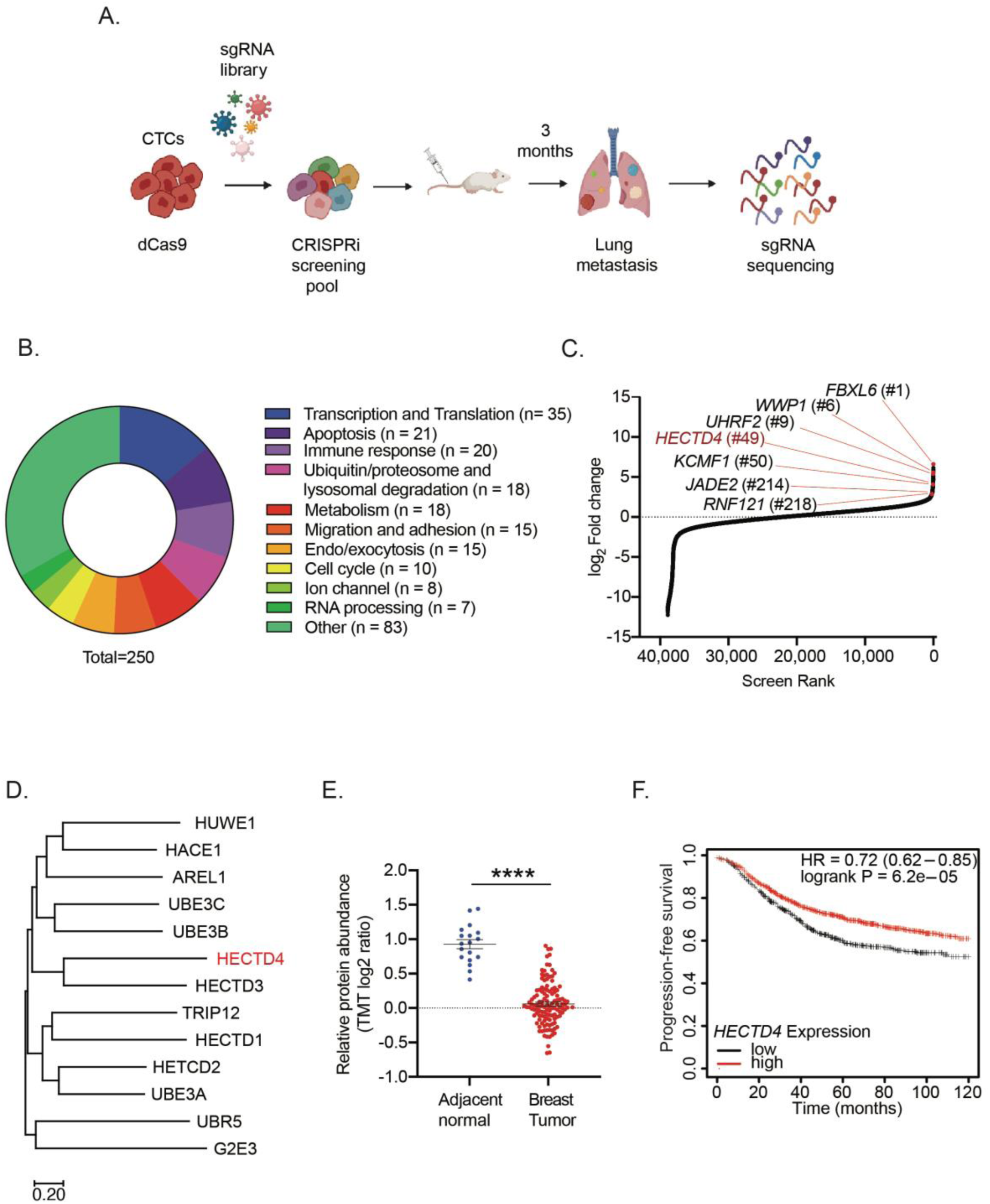
Suppression of CTC-mediated metastasis by HECTD4 in a CRISPR-i screen. **(A)** Schematic diagram of the *in vivo* CRISPR inactivation screen in CTCs. **(B)** Classification of known functions of the top 250 genes identified in the *in vivo* CRISPR inactivation (CRISPRi) screen. **(C)** Distribution of ranked screen scores, according to the fold change compared to the input (log2 FC). The top genes classified as E3 ubiquitin ligases are indicated with *HECTD4* highlighted in red. **(D)** The evolutionary tree showing the 13 members of the HECT subfamily. The distances are in the units of the number of amino acid substitutions per site. **(E)** Proteomics data from breast tumor tissues showing a reduction in HECTD4 protein in tumors compared with normal breast tissue. Significance was calculated using unpaired student t-test. **(F)** Kaplan-Meier plot shows that breast cancer patients with high *HECTD4* expression (n=1433) in their tumors have improved progression-free survival compared with those with low *HECTD4* expression. Significance determined by log-rank (Mantel-Cox) test. (* p<0.05, ** p<0.01, *** p<0.001, **** p<0.0001)

Among the three major gene families of E3 ubiquitin ligases, the HECT family encompasses 13 proteins sharing a <u>H</u>omologous to <u>E</u>6AP <u>C</u>-<u>T</u>erminus (HECT) domain, which catalyzes the transfer of the ubiquitin moiety onto target proteins (13). *HECTD4* shares a catalytic HECT domain with 12 other members of this family of E3 ubiquitin ligases (**FIG. 1D**). HECT family members have known targets implicated in breast cancer development and progression (14), but HECTD4 has remained uncharacterized, with no known function or other recognizable functional domain and noteworthy for its very large size (460 kD). Inherited genetic variants of *HECTD4* are associated with neurological disease and increased risk for diabetes and hypertension (13).

To support a potential role in cancer, we first explored clinical datasets, observing that both *HECTD4* mRNA and protein expression are considerably reduced in primary and metastatic breast cancers, compared with normal breast tissue (**FIG. S1A, 1E**) (15). Moreover, lower *HECTD4* expression in breast cancer is associated with reduced progression-free survival (**FIG. 1F**) (16). Among histological subsets of breast cancer, *HECTD4* expression is lowest in the most aggressive subtype, triple-negative breast cancer (TNBC), a proportion of which display allelic losses around the gene locus (**FIG. S1B, S1C**). In addition to breast cancer, *HECTD4* expression is reduced in multiple other tumor types compared with corresponding normal tissues, including glioblastoma and esophageal cancers (**FIG. S1D, S1E**). We therefore selected *HECTD4* as the CRISPR-i hit for detailed analysis.

### Anchorage-independent proliferation and increased tumorigenesis following HECTD4 depletion

To validate the effect of *HECTD4* suppression, we turned to the TNBC cell line MDA-MB-231, which is highly invasive and metastatic, as well as genetically manipulable. Despite their high baseline tumorigenesis, MDA-MB-231 cells with shRNA-mediated *HECTD4* knockdown (>80%) doubled in tumor size in immunosuppressed NSG mice, compared with sh-scramble controls (457.1 µg ± 118.1 vs 204.9 µg ± 51.01, p = 0.0078; **FIG. 2A, S2A**). To futher quantify this effect, we performed chimeric tumorigenicity experiments by mixing GFP- and mCherry-tagged tumor cells (**FIG. 2B UPPER, S2B**). In a control experiment, a 1:1 mixture of GFP- and mCherry-tagged MDA-MB-231 cells (GFP-shControl; *HECTD4*-WT vs GFP-mCherry-shControl; *HECTD4*-WT) (n=4-5 per group) shows persistence of the 1:1 ratio in primary tumor (orthotopic mammary fat pad), comparable with the ratio in the day 0 input cell population, as measured by FACS analysis. Following resection of the primary tumor (day 36 post-inoculation) to allow for the development of metastases (day 62), the mCherry/GFP ratio remains constant in lung and liver metastases. In marked contrast, inoculation of a 1:1 mixture of control and *HECTD4*-KD cells (GFP- shControl; *HECTD4*-WT vs GFP-mCherry-*HECTD4*-KD; *HECTD4*-KD) shows a dramatically increased representation of the *HECTD4*-deficient cells in both primary and metastatic lesions. In all primary tumors, *HECTD4*-KD cells constituted >80% of cells, and as high as >95% in three of five tumors. Similarly, *HECTD4*-KD cells contributed to >70% of metastatic tumor cells in the lungs and >90% in the liver (**FIG. 2B LOWER, S2C, S2D**). Thus, even in the highly invasive MDA-MB-231 breast cancer cells, suppression of *HECTD4* further enhances both primary and metastatic tumorigenesis.

**Figure 2.**
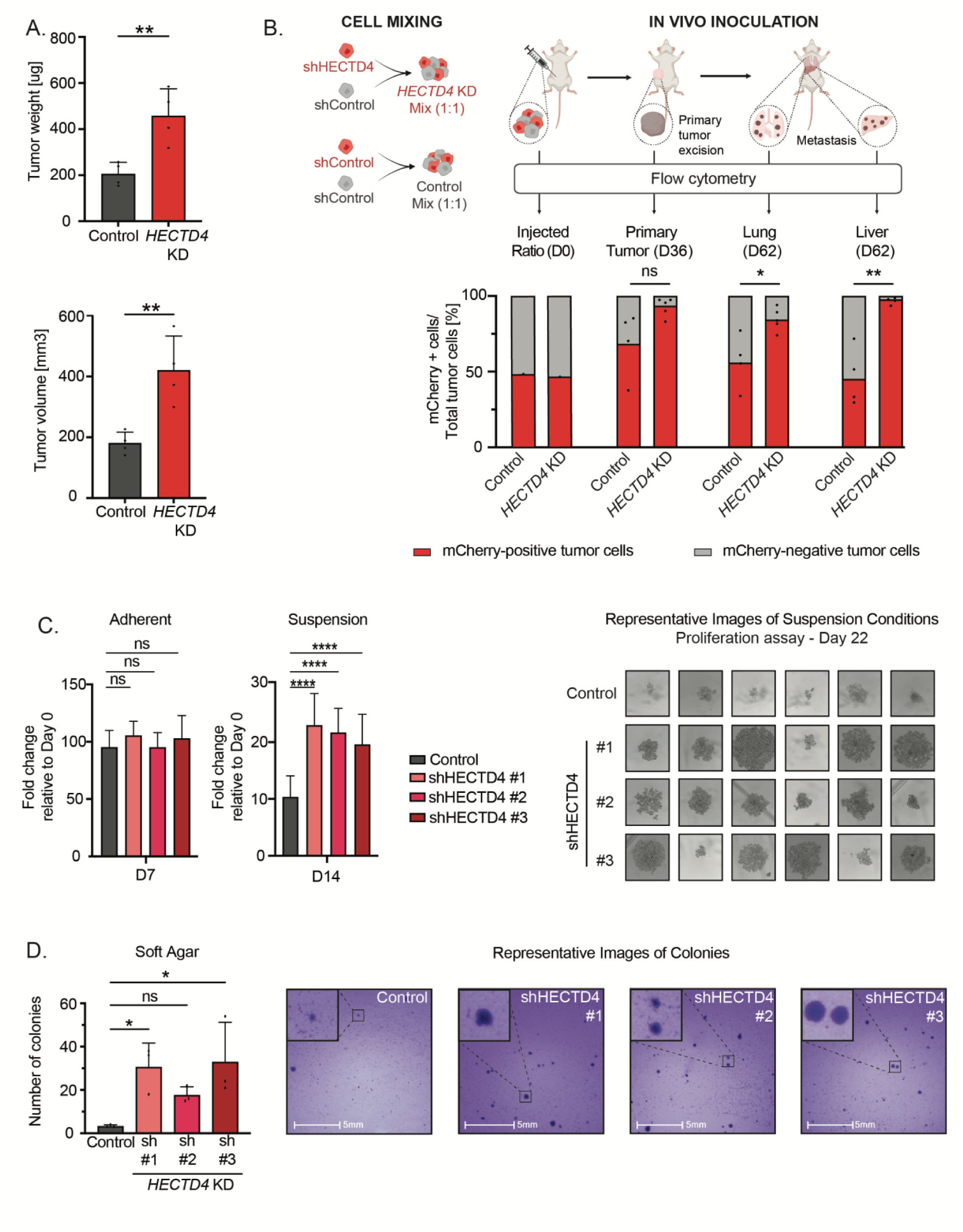
Anchorage-independent proliferation and increased tumorigenesis following *HECTD4* depletion. **(A)** The initial *in vivo* validation experiment comparing *HECTD4*-KD (sh #1) cells with scrambled control cells. MDA-MB-231 cells with *HECTD4*-KD (sh #1) or scrambled control were injected separately into the mammary fat pad of NSG mice (n= 4 mice for the control group, n=4 mice in *HECTD4*-KD group). Primary tumors were harvested after 34 days and assessed for tumor weight and tumor volume. Bar graph shows tumor weights **(Upper)** and volumes **(Lower)** measured at day 34. Error bars represent mean +/- SD. Significance was calculated using unpaired student t-test. The mRNA levels of *HECTD4* in the cells used for this experiment are shown in **Fig. S2A. (B)** Schematic representation of the tumor cells mixing experiment to interrogate the effects of HECTD4 on tumorigenesis. shControl cells (GFP-shControl; *HECTD4*-WT, shown in gray) were mixed with either GFP-tagged and mCherry labelled *HECTD4*-knockdown cells [GFP-mCherry-*HECTD4*-KD, shown in red] (experimental cohort) or with GFP-tagged and mCherry labelled shControl cells [GFP-mCherry-shControl; *HECTD4*-WT, shown in red] (control cohort). A 1:1 mixture of GFP-shControl and GFP-mCherry-*HECTD4*-KD or GFP-shControl and GFP-mCherry-shControl was inoculated into the mammary fat pad of immunodeficient NSG mice (n= 4 mice for control mixture; n = 5 mice for KD mixture, see Methods for more detail). The mixtures were analyzed by flow cytometry right before inoculation to ensure that each of the populations were equally represented in the mixture inoculated into the mammary fat pad at day 0. The primary tumors were resected after 36 days via survival surgery of the mice and the colonized organs – the lungs and livers – were harvested after 62 days. The ratio of the GFP: mCherry populations was analyzed by flow cytometry in the control and experimental cohorts **(Upper)**. The bars represent the percentage of green (GFP-shControl, shown in gray) and red (GFP-mCherry-shControl or GFP-mCherry-*HECTD4*-KD, shown in red) cells in the primary and metastatic tumors. Significance was calculated with unpaired student t-test, the dots indicate the mCherry positive samples **(Lower)** The mRNA levels of *HECTD4* in the cells used for this experiment are shown in **Fig. S2B.** *In vitro* growth of *HECTD4*-depleted MDA-MB-231 cells compared to scrambled control cells under adherent 2D-culture on day 7. Error bars represent mean +/- SD. Significance was calculated using two-way ANOVA, Tukey’s multiple comparison test (day 7) **(Left)**. *In vitro* growth of *HECTD4*-depleted MDA-MB-231 cells compared to scrambled control cells under anchorage-independent suspension conditions (ultra-low adherent plates) on day 14. Error bars represent mean +/- SD. Significance was calculated using the two-way ANOVA, Tukey’s multiple comparison test (day 14) **(Middle)**. Representative photomicrographs of MDA-MB-231 cells grown in anchorage-independent suspension conditions (ultra-low adherent plates) on day 22 **(Right)**. The mRNA levels of *HECTD4* in the cells used for this experiment are shown in **Fig. S2A. (D)** Single MDA-MB-231 cells with either *HECTD4* depletion or scrambled control were seeded on soft-agar coated 6-well plates (10,000 cells/well). After 2 weeks, the colonies were stained with crystal violet, imaged and the number of colonies was counted using the HALO software. Error bars represent mean +/- SD. Significance was calculated using one-way ANOVA test, with Dunnett’s multiple comparison test **(Left)**. Representative images of these colonies **(Right)** The mRNA levels of *HECTD4* in the cells used for this experiment are shown in **Fig. S2F**. (* p<0.05, ** p<0.01, *** p<0.001, **** p<0.0001)

Despite the striking effect of *HECTD4* knockdown *in vivo*, it has no effect on *in vitro* proliferation of MDA-MB-231 cells under standard adherent 2D culture conditions (**FIG. 2C LEFT, S2A**). Remarkably, however, *HECTD4*-KD dramatically promotes anchorage-independent growth of these cells upon plating onto low adherence plates, where they form large spheroid-like structures, that are not present in *HECTD4*-expressing control cells (**FIG. 2C MIDDLE, RIGHT**). *HECTD4*-KD cells grown in suspension show increased PCNA expression, consistent with their increased proliferation (**FIG. S2E**). The increased anchorage-independent proliferation in suspension by *HECTD4*-KD cells is also evident by increased colony formation in soft agar assays, a commonly used hallmark of tumorigenenesis (**FIG. 2D LEFT, RIGHT, S2F**). Taken together, these findings show that depletion of *HECTD4* results in enhanced anchorage-independent proliferation *in vitro* and increased tumorigenesis *in vivo*.

### Regulation of COX-2 by HECTD4

To first demonstrate that HECTD4 has ubiquitin ligase activity, we used CRISPR prime-editing to engineer an ALFA tag onto the N-terminus of endogenous HECTD4, enabling efficient pull-down of native protein using a tag-specific nanobody (**FIG. S3A, S3B**). We then performed an auto-ubiquitination assay *in vitro* by adding essential components of the ubiquitin cascade, including E1, E2, ubiquitin and ATP to the purified HECTD4. Western blot analysis of the products shows that the addition of HECTD4 results in the formation of polyubiquitin conjugates, which is characterized by the presence of higher-order molecular weight species of additional ubiquitin moieties. Addition of the de-ubiquitinating enzyme Ubiquitin Specific Peptidase 2 (USP2) results in the loss of this ubiquitin smear (**FIG. S3C, 3A**). The HECT domain of HECTD4 contains a highly conserved cysteine at its C-terminal end, which, in other family members, is required for ubiquitin-binding and transfer from E2 to the target protein. Accordingly, we cloned a “short-active” form of HECTD4, encoding the ALFA-tagged isolated HECT domain, and mutated the active site cysteine C4396 to alanine to generate a “short-mutant” construct (**FIG. 3B UPPER**). Transfection of the “short-active”, but not the “short-mutant” construct, into MDA-MB-231 cells shows auto-ubiquitination, following immunoprecipitation from cellular extracts (**FIG. 3B LOWER, S3D**).

**Figure 3.**
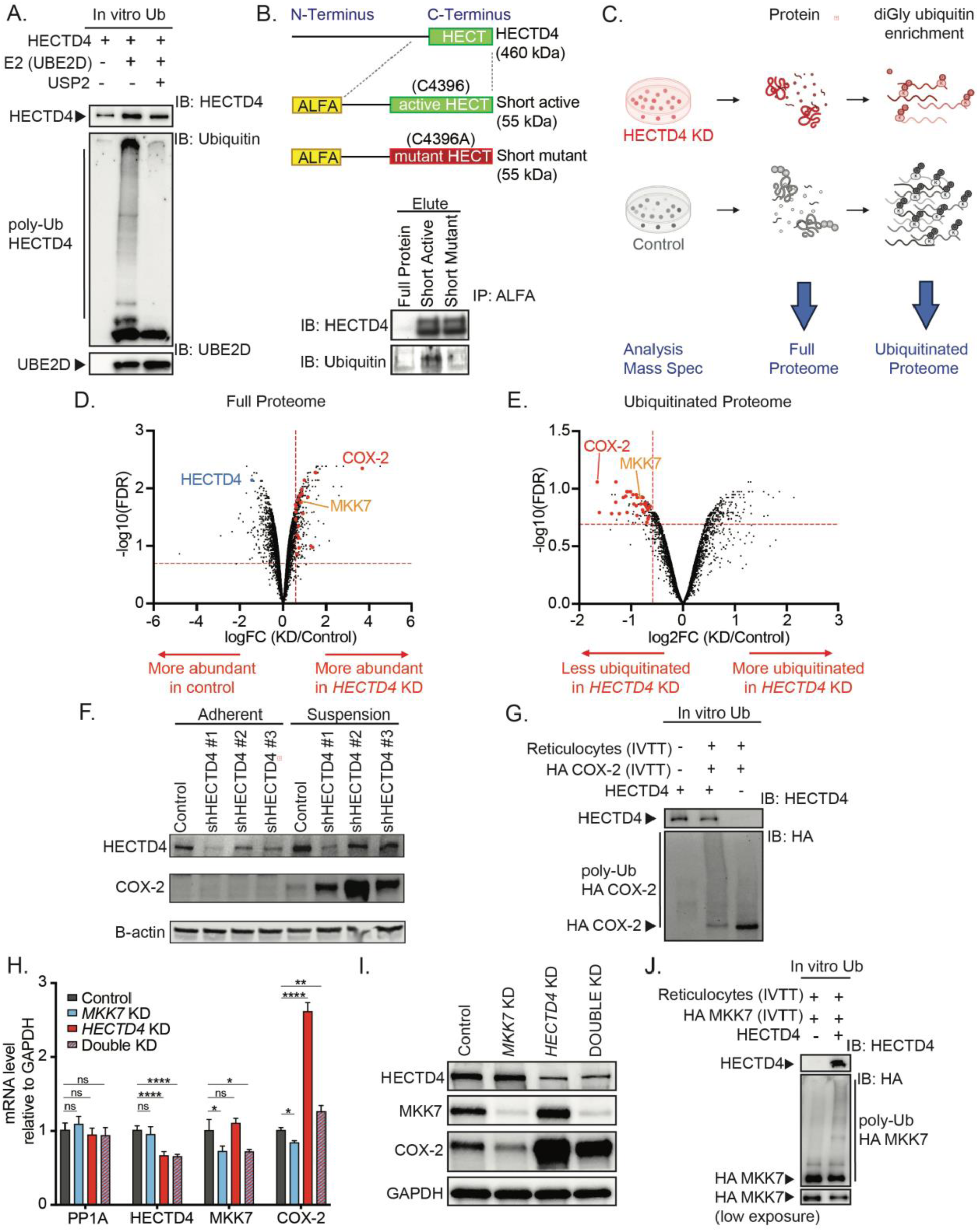
HECTD4 modulates the stability of COX-2 and its regulatory kinase MKK7. **(A)** HECTD4 catalyzes ubiquitin chain formation *in vitro*. Active HECTD4 was immunoprecipitated from freshly lysed cells expressing the full-length alfa-tagged HECTD4 using a nanobody against the alfa-tag. Addition of E1, UBE2D, ATP and ubiquitin results in the formation of polyubiquitin conjugates. The addition of the de-ubiquitinating enzyme Ubiquitin Specific Peptidase 2 (USP2) leads to the loss of the polyubiquitin chain. The potential reaction products were detected by western blotting against HECTD4, ubiquitin and UBE2D. **(B)** Schematic representation of the cloning strategy of the C-terminal end of HECTD4. The “short active” product contains the reference sequence of HECT and 380 nucleotides upstream, including the putative catalytic cysteine, C4396, in green. In the “short mutant” form, this cysteine was mutated into an alanine (C4396A). To facilitate the immunoprecipitation and increase its specificity, both products were tagged at the N-terminus with an ALFA tag. The resultant protein has an expected molecular weight of 55 kDa **(Upper)**. Lysates from cells transfected with the short-active and short-mutant forms of alfa-HECTD4 were flowed through a ALFA-tag activated column. The input lysate, flow-through, and the eluate were blotted and probed with antibodies against HECTD4 and ubiquitin. Untransfected parental cells are shown as control. The elute is shown here; the input lysate and flow-through are shown in **Fig. S3D.** Ubiquitin blotting detects signal from the short active form of HECTD4 but not from the short mutant in which C4396 is converted to Alanine **(Lower)**. **(C)** Schematic representation of the quantitative proteomics experiment to identify HECTD4 substrates. *HECTD4*-KD and control cells were cultured under anchorage-independent conditions for four days. Cells were lysed in a urea-based buffer, followed by protease-digestion of the proteins. A fraction of the complete lysate was conserved for analysis by full proteome mapping. The remaining peptides were purified by reversed-phase, solid-phase extraction. Using an antibody targeting the diGly ubiquitin remnant motif (K-ε-GG), ubiquitinated peptides were then captured by immunoprecipitation and eluted, to concentrate them for LC-MS/MS analysis. **(D)** Volcano plot of the complete proteome of *HECTD4*-KD MDA-MB-231 cells versus control cells. The x-axis shows the log2 FC of each identified protein (6266), and the y-axis shows the corresponding –log10 P adjusted value. The HECTD4 protein is marked in blue; enriched proteins in the *HECTD4*-KD cells are on the right side of the graph with a positive log2FC. 25 proteins highlighted in red were found to accumulate in the *HECTD4*-KD cells (FC >1.5 and FDR < 0.25, thresholds in dotted lines) and to have their ubiquitinated forms less abundant in the same cells (FC < -1.5 and FDR < 0.25, thresholds in dotted lines) in the diGly screen. COX-2 ranks as the second most enriched protein (>12-fold increase). **(E)** Volcano plot of the ubiquitinated peptides captured by immunoprecipitation from *HECTD4*-KD cells versus control cells. The x-axis shows the log2 FC of each identified peptides (3311) aligning to 1453 proteins, and the y-axis shows the corresponding –log10 P adjusted value. HECTD4 substrates are expected to be less ubiquitinated in the *HECTD4*-KD cells, thus exhibit peptides on the left side of the graph with a negative log2FC. The 25 proteins described in Fig. 3D. are also highlighted in red. COX-2 is the protein with the most reduced ubiquitination (>3-fold reduction compared to normal cells). **(F)** Western blot of MDA-MB-231 cultured in regular adherent conditions (adherent) and placed in suspension (suspension) showing the weak expression of COX-2 in adherent cells compared to suspension cells. *HECTD4*-KD cells express high levels of COX-2 compared with control. HECTD4 protein expression is also upregulated in suspension. **(G)** HECTD4 ubiquitinates COX-2 *in vitro*. Potential reaction products were detected by western blotting against HECTD4 and HA-tag. The smear demonstrates the presence of a poly-ubiquitinated HA-COX-2. **(H)** qRT-PCR of *PP1A, HECTD4, MKK7* and *COX-2* mRNA upon depletion of *HECTD4* only (mix of short hairpins targeting HECTD4), of *MKK7* only (mix of 4 siRNA targeting MKK7, shown in blue) and of a combined KD (mix of shHECTD4 and mix of siMKK7, shown in red and blue stripes). mRNA collected 3 days after siRNA transfection from cells growing in suspension. *COX-2* mRNA levels increase upon *HECTD4* depletion, and *MKK7*-KD reverses it. Error bars represent mean +/- SD. Significance was calculated using one-way ANOVA test, with Dunnett’s multiple comparison test. **(I)** Western blot of a similar experiment described in Fig. 3H, with protein lysate harvested after 4 days. **(J)** HECTD4 also ubiquitinates MKK7 *in vitro*. Potential reaction products were detected by western blotting against HECTD4 and HA-tag. The smear demonstrates the presence of a poly-ubiquitinated HA-MKK7. Lower exposure of the same gel shown below. (* p<0.05, ** p<0.01, *** p<0.001, **** p<0.0001)

To identify proteins targeted for degradation by HECTD4, we conducted two complementary proteomic screens, comparing *HECTD4*-KD cells versus controls. In both screens, MDA-MB-231 cells were cultured under low adherence conditions (**FIG. 3C**). First, we performed quantitative mass spectrometry-based proteomics (17) on total cellular lysate followed by tryptic digest to identify proteins with increased concentration following *HECTD4* knockdown (**FIG. 3D**). Second, we enriched for tryptic peptides containing lysine residues that carry a diGly remnant of ubiquitin (18) at lysine residues and we used mass spectrometry to identify proteins with decreased ubiquitination upon *HECTD4* knockdown (**FIG. 3E**). Looking at the intersection of these two orthogonal assays, a total of 25 proteins satisfied our protein concentration change filter criteria (FC (KD/Control) >2 at an FDR < 0.25). Of these proteins, the cyclooxygenase protein COX-2 was >12-fold more abundant and showed the most pronounced decrease of ubiquitination (>3-fold) upon *HECTD4* knockdown (**FIG. 3D, 3E**).

Western blotting confirms a dramatic increase in COX-2 protein levels in *HECTD4*-KD cells compared to control in MDA-MB-231 cells (**FIG. S3E, S3F**). Remarkably, this effect is only evident when MDA-MB-231 cells are maintained under anchorage-independent conditions (**FIG. 3F**). COX-2 levels in MDA-MB-231 cells are undetectable under standard 2D culture conditions, but COX-2 expression becomes detectable when cells are placed in suspension; *HECTD4*-KD mediates a significant increase in COX-2 levels under suspension, but not in 2D culture conditions (**FIG. 3F**). Endogenous HECTD4 levels also increase upon anchorage-independent growth, raising the possibility that it may play a physiological role in attenuating the increased COX-2 expression under stressful conditions. The rise in COX-2 protein levels in the absence of *HECTD4* under suspension conditions is also seen in three other breast cancer cell lines HCC1143, BT-549 and MDA-MB-468 (**FIG. S3G, S3H, S3I** ). To further validate the inverse correlation between HECTD4 and COX-2 proteins, we analyzed a previously published proteomic dataset from 21 different human breast cancer cell lines (19), which confirmed the negative correlation between HECTD4 and COX-2 protein levels (N = 21, r = -0.426, p = 0.0541; **FIG. S3J**). Finally, to confirm the direct ubiquitination of COX-2 by native HECTD4, we purified ALFA-tagged endogenous HECTD4 from cells in suspension culture and found that purified HECTD4 can ubiquitinate recombinant HA-tagged COX-2 using in vitro ubiquitination assays. Western blotting demonstrates a smear of poly-ubiquitinated HA-COX-2, coincident with reduction of the unmodified HA-COX-2 (**FIG. S3K, 3G**).

In addition to the dramatic increase in COX-2 protein levels in *HECTD4*-KD cells grown in suspension, we also observed a modest increase in *COX-2* mRNA under these conditions (**FIG. S4A**). Two of the 25 candidate HECTD4 target proteins identified in our proteomic screen (**FIG. 3D, 3E**), MEK1 (MAP2K1) and MKK7 (MAP2K7) (20, 21), are mitogen-activated kinase (MAPK) family members known to activate transcription factors c-Jun, ATF2, and Elk1, which function to induce *COX-2* gene expression (**FIG. S4B**). In *HECTD4*-KD cells, the MEK1 inhibitor trametinib does not alter *COX-2* mRNA or protein levels, despite effectively inhibiting its direct target ERK1/2 (**FIG. S4C, S4D, S4E**). By contrast, either short-interfering RNA (siRNA)-mediated suppression of MKK7 or treatment with the MKK7 inhibitor 5Z-7-oxozeaenol (22) reverses the *HECTD4*-KD-dependent increase in *COX-2* mRNA (**FIG. 3H, S4F**). Importantly, in *HECTD4-MKK7* double KD cells, COX-2 protein remains elevated, pointing to direct protein stabilization by loss of HECTD4 **(FIG. 3I**). To confirm that MKK7 itself is a HECTD4 target protein, we demonstrated its accumulation at the protein level (**FIG. S4G**) in *HECTD4*-KD cells, without changes in *MKK7* mRNA (**FIG. S4H**). ALFA-tag pull-down of endogenous HECTD4 also mediates ubiquitination of *in vitro* translated HA-tagged MKK7 (**FIG. S3K, 3J**).

Thus, HECTD4-mediated ubiquitination regulates COX-2 expression both directly and indirectly, through ubiquitin-mediated degradation of COX-2 protein itself and through targeting of its upstream transcriptional regulator MKK7, respectively.

### HECTD4 modulation of COX-2-dependent tumorigenesis and metastasis

Given the dramatically increased COX-2 protein expression upon *HECTD4* depletion in epithelial cells that lose matrix attachment, we sought to determine whether the tumorigenic phenotype evident in *HECTD4*-KD cells is dependent upon COX-2. Indeed, shRNA-mediated suppression of COX-2 abrogates the increased proliferation of *HECTD4*-KD cells when these are grown under anchorage-independent conditions (**FIG. 4A, S5A, S5B, S5C**) as does the selective COX-2 inhibitor, celecoxib (**FIG. 4B, S4C**). Remarkably, we find that soft agar colony formation itself is dependent upon COX-2 activity, suggesting an essential contribution of COX-2 for cell proliferation under anchorage-independent states. Consistent with its upstream regulation of COX-2, *HECTD4*-KD fails to rescue the *COX-2*-KD phenotype (**FIG. 4C LEFT, RIGHT, S5D**).

**Figure 4.**
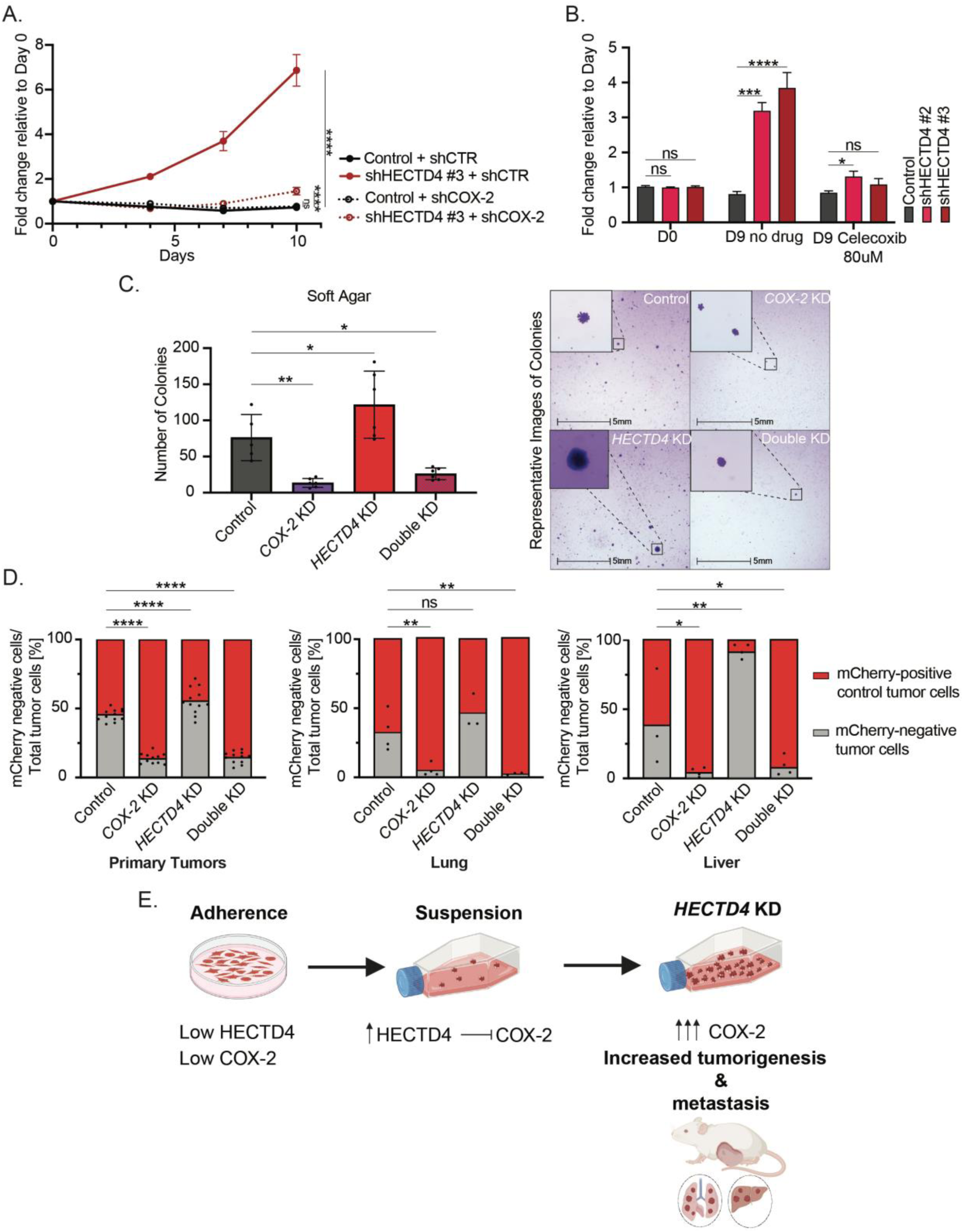
HECTD4 modulation of COX-2-dependent tumorigenesis and metastasis. **(A)** Depletion of *HECTD4* increases cell proliferation under anchorage-independent, suspension conditions (ultra-low adherent culture dish) compared to scrambled control. Depletion of *COX-2* in the *HECTD4*-depleted cells reverts this phenotype. Cell viability and proliferation measured by CellTiter Glo luminescence. Error bars represent mean +/- SD. Significance was calculated using with two-way ANOVA test, with Tukey’s multiple comparison test (Day 10). The mRNA levels of *PP1A, HECTD4* and *COX-2* in the cells used for this experiment are shown in **Fig. S5A.** Only the cells harboring the control and shHECTD4 #3 shRNA’s are shown in this figure. Cells with the control and shHECTD4 #1 shRNA’s are shown in **Fig. S5B. (B)** *HECTD4*-KD cell proliferation is increased under anchorage-independent, suspension conditions (ultra-low adherent culture dish) compared to scrambled control which remain stable after 9 days without treatment. Addition of celecoxib (80 umol/L) reverses the proliferation advantage of *HECTD4*-KD cells. Cell viability measured by CellTiter Glo luminescence upon celecoxib treatment. Error bars represent mean +/- SD. Significance was calculated using one-way ANOVA test, with Dunnett’s multiple comparison test. The baseline mRNA levels of *PP1A, HECTD4* and *COX-2* in the cells used for this experiment are shown in **Fig. S4C.** The same cells were used in the experiments in **Fig. S4D. (C)** Four different groups (scramble control, *COX-2*-KD, *HECTD4*-KD and double KD) of single MDA-MB-231 cells were seeded on 6-well plates coated with soft agar (10,000 cells/well). After 2 weeks, the colonies were stained with crystal violet and counted using the HALO software. Error bars represent mean +/- SD. Significance was calculated using one-way ANOVA test, with Dunnett’s multiple comparison test **(Left)**. Representative images of colonies from this experiment **(Right).** The mRNA levels of *HECTD4* and *COX-2* in the cells used for this experiment are shown in **Fig. S5D.** The same cells were used in the *in vivo* mixing experiment in Fig. 4D. **(D)** The fraction of the four GFP+/mCherry-conditions i.GFP-shControl, ii.GFP-*COX-2*-KD, iii.GFP-*HECTD4*-KD, iv.GFP-Double-KD [%] (n = 6 mice per group, primary tumors were bilaterally injected into the mammary fat pads of each mice, see Methods for more detail) in the tumor arising from the mixed tumor cell populations were calculated by flow cytometry, as depicted in the schematic **Fig. S6A**. The bars represent the percentage of red (GFP-mCherry-shControl, shown in red) and green (i.GFP-shControl, ii.GFP-*COX-2*-KD, iii.GFP-*HECTD4*-KD, iv.GFP-Double-KD, shown in gray) cells in the primary **(Left)** and metastatic tumors **(Middle-Lungs, Right-Liver)**. Significance was calculated using one-way ANOVA test, with Dunnett’s multiple comparison test, the dots indicate the mCherry negative samples. The mRNA levels of *HECTD4* and *COX-2* in the cells used for this experiment are shown in **Fig. S5D.** The same cells were used in the soft agar experiment in Fig. 4C. In vitro, day 0 injection ratios of the cells used in this experiment are shown in **Fig. S6B**. The volumes of the primary tumors are shown in **Fig. S6C. (E)** The diagram of the proposed model showing HECTD4 acting as a tumor and metastasis suppressor, responsible for the regulation of COX-2. HECTD4 and COX-2 levels are undetectable in adherent state. The levels of both proteins rise in anchorage-independent conditions and *HECTD4*-KD results in massive amounts of COX-2 accumulation. In anchorage-independent state, *HECTD4*-KD with high COX-2 levels leads to increased growth *in vitro* as well as increased tumorigenesis and metastasis *in vivo*. (* p<0.05, ** p<0.01, *** p<0.001, **** p<0.0001)

To test whether the *in vivo* tumorigenic phenotype of *HECTD4*-KD MDA-MB-231 cells is also dependent on COX-2, we set up a cell mixing experiment analogous to that described above (**FIG. 2B, S6A**). Using pair-wise 1:1 mixing experiments with control cells, we tested the tumorigenic contributions of *COX-2*-KD, *HECTD4*-KD and the double *HECTD4/COX-2*-KD, using tagged GFP and mCherry markers (**FIG. S5D, S6A-C**). As with our previous mixing experiment, primary tumors were harvested at day 25 after orthotopic inoculation, and lungs and liver were collected at day 54 to evaluate metastatic growth following removal of the primary tumor. FACS analysis of primary tumors and metastasis confirms the increased tumorigenesis of *HECTD4*-KD cells (**FIG. 4D**). *COX-2*-KD cells are profoundly depleted from tumors, consistent with the tumor suppressing effect of COX-2 inhibition (23, 24). In the double KD, the simultaneous suppression of COX-2 completely abrogates the increased tumorigenic effect of *HECTD4*-KD (**FIG. 4D LEFT**). Similar results are observed in the metastatic lung and liver lesions, where *COX-2*-KD alone dramatically suppresses metastatic dissemination and the double KD completely reverses the growth advantage conferred by the *HECTD4*-KD (**FIG. 4D MIDDLE, RIGHT**). Taken together, the tumor and metastasis suppressor effect of HECTD4 is dependent on suppressing COX-2 activity.

## Discussion

We demonstrate a tumor and metastasis suppressor role for HECTD4, a previously uncharacterized member of the HECT family of E3 ubiquitin ligases. Our initial CRISPR-i screen scored HECTD4 as a suppressor of metastasis initiation by breast cancer patient-derived CTCs, and our follow-up analyses point to a critical effect in anchorage-independent cell proliferation, with COX-2 as a major degradation target. Together, these observations suggest a potentially critical role for this prostaglandin-synthetic pathway in the survival of epithelial cancer cells in the circulation (**FIG. 4E**).

Among the best-studied protein degradation pathways implicated in tumorigenesis are von Hippel Lindau (VHL) targeting of HIF, MDM2 degradation of TP53, FBXW7 destabilization of MYC and SKP2 degradation of p27/CDKN1B (25). Furthermore, several different deubiquitinases have also played a role in cancer, such as USP11 in the regulation of the estrogen receptor in breast cancer, USP15 in TGF-β signaling in glioblastoma and USP26 in androgen receptor regulation (26–29). Our initial loss-of-function *in vivo* CRISPR-i screen identified previously characterized high-ranking hits in the ubiquitin-proteasomal and lysosomal degradation pathways, *FBXL6, WWP1, UHRF2, KCMF1, JADE2 and RNF121.* However, HECTD4 had remained uncharacterized, with no known function or substrate, leading us to further pursue its role in tumorigenesis and metastasis. Interestingly, within the HECT gene family, not all proteins mediate proteolytic ubiquitination. HERC2 promotes BRCA1 degradation and WWP1 promotes ubiquitination and degradation of p27, ErbB4 and other substrates, but SMURF1-mediated ubiquitination stabilizes estrogen receptor α, while HECTD3 has opposing ubiquitination effects, degrading caspase 8, while stabilizing the NFkB regulator MATL1 (13). Our studies of the previously uncharacterized HECT gene family member HECTD4 thus establish its catalytic functions as a degradative E3 ubiquitin ligase, with COX-2 as a major degradation target. We note that we previously conducted a CRISPR-activation (CRISPR-a) screen in breast cancer CTCs, nominating multiple regulators of protein translation, including core structural components of the ribosome, as enhancers of CTC-mediated tumorigenesis (4). Together, these loss of function and gain of function CRISPR screens highlight the relevance of protein synthesis and degradation as regulatory factors in tumorigenesis and metastasis.

A role for the ubiquitin ligase HECTD4 in cancer is supported by clinical databases indicating a correlation between reduced HECTD4 expression in breast cancer and an adverse progression-free survival (16). The most striking characteristic, drawn from our experimental models, is its unique contribution to anchorage-independent proliferation. Indeed, HECTD4 is expressed at low levels in cultured cells grown under standard 2D matrix attachment conditions, but it is induced as these cells lose their anchorage dependence. It is only under anchorage-independent culture conditions that *HECTD4* knockdown mediates an effect on cell proliferation. Anoikis is thought to be the predominant form of cell death facing epithelial cancer cells that intravasate into the bloodstream, and its suppression is likely to contribute to the ability of CTCs to survive in circulation and ultimately give rise to metastatic lesions. In this context, the targeting of COX-2 by HECTD4, both through direct protein ubiquitination and through suppression of its regulatory kinase MKK7, highlights a potentially important role for this prostaglandin synthetic pathway in the survival of metastatic precursors in circulation.

COX-2 is the rate-limiting synthetic enzyme in the synthesis of the prostaglandin PGE2, mediating an array of pro-tumorigenic inflammatory signals (5). These include secretion of growth factors mediating angiogenesis, cell proliferation, apoptotic resistance, and cell invasion. The role of COX-2 in enhancing cancer has been primarily derived from seminal epidemiological studies of chemo-prevention through administration of non-steroidal anti-inflammatory drugs (NSAID) (7, 30, 31). These studies demonstrated a potent suppression of cancer initiation across colorectal and other cancer types. However, the broad utility of NSAIDs and selective COX-2 inhibitors as chemo-preventive agents is negated by their concurrent activation of thrombotic pathways, resulting in increased cardiovascular events in individuals who do not have cancer (7, 32). Nonetheless, additional studies have also demonstrated a beneficial effect of COX-2 inhibition in the setting of metastatic progression (9), which may present a different risk/benefit calculation, depending on the relative risk of cancer recurrence versus cardiovascular complication.

In multiple mouse models, COX-2 inhibition prevents or reduces the development of metastasis from breast, colorectal and lung tumors (33, 34). While clinical studies of metastatic suppression are challenging to undertake, one meta-analysis indicates that daily aspirin use is associated with a reduced risk of presenting with metastatic adenocarcinoma, or of developing metastases following treatment of a primary cancer (10). In a recent study, including celecoxib as an adjuvant to treat stage III colon cancer was associated with improved disease-free survival and overall survival (35). However, another study found no effect of high-dose aspirin in preventing early recurrence of breast cancer (11). Of note, the combination of short-term COX-2 inhibition with chemotherapy or other modalities for the treatment of established metastatic disease has proven ineffective. Taken all together, translating the striking effect of COX-2 inhibition in preventing cancer metastasis in mouse models to human clinical studies will require careful consideration of metastatic risk stratification, as well as drug dosage, duration of treatment and appropriate endpoints. In this context, the role of COX-2 inhibition in the prevention of blood-borne metastasis may impact the optimal design of clinical studies.

Our observations point to a distinct mechanism whereby COX-2 may mediate its pro-metastatic effects. While most studies have focused on the inflammatory output of prostaglandins in cancer initiation, the specific anchorage-independent growth context in which HECTD4 regulates COX-2 expression indicates a distinct role in sustaining epithelial cancer cells as they circulate in the bloodstream. Consistent with such a cell autonomous and non-inflammation mediated mechanism, COX-2 has been reported to enhance expression of E-cadherin under 3D culture conditions, thereby sustaining reattachment of suspended cells onto matrix and enhancing tumorigenesis (36). Thus, suppressing the ability of circulating cancer cells to survive in the bloodstream and initiate metastatic growth through the HECTD4-COX-2 axis reported here highlights a novel therapeutic opportunity, which may warrant clinical investigation of metastasis prevention in carefully selected clinical contexts.

## Materials and Methods

### Cell culture

For the initial CRISPR screen, the CTCs Brx-142 were grown in suspension in anchorage-independent, ultra-low attachment plates (Corning) in tumor sphere media, consisting of RPMI-1640 with GlutaMAX (Gibco) supplemented with EGF (20ng/mL), FGF (20ng/mL), 1X B27, and 1X antibiotic/antimycotic (Life Technologies), in 4% O2, as previously described (3, 37). They were authenticated by RNA-seq and DNA-seq.

Human mammary cancer cell lines MDA-MB-231, HCC1143, BT-549 and MDA-MB-468 cells were all obtained from ATCC, derived from a female patient. Cell cultures were maintained on adherent plates (Corning) at 37 °C and 5% CO2 in humidified culture incubators. MDA-MB-231 and MDA-MB-468 cells were grown in DMEM high glucose medium (Gibco) with 10% FBS (Gibco) and 1X Pen/Strep (Gibco). HCC1143 and BT-549 cells were grown in RPMI-1640 with GlutaMAX (Gibco) supplemented with 10% FBS (Gibco) and 1X Pen/Strep (Gibco). To assess HECTD4, COX-2 and MKK7 levels in suspension, cells were seeded in anchorage-independent, ultra-low attachment plates (Corning) and cultured for 3 to 5 days prior to harvest. The cell lines used in this study were checked for mycoplasma every 2 months using the Mycoalert kit (Lonza).

### Lentiviral production and infection of the CTCs with the CRISPR inactivation library

HEK293T cells were grown in high-glucose DMEM supplemented with 10% fetal bovine serum and 1% penicillin/streptomycin. For the CRISPR inactivation library production, cells were transfected at ∼80% confluency in T225 flasks. For each flask, 15.3µg pMD2.G (Addgene #12259), 23.4µg psPAX2 (Addgene #12260), and 30.6µg pooled library plasmid were transfected using 270µL Lipofectamine 2000 (Invitrogen #11668019) and 297µL PLUS reagent, as previously described (4). 24h after transfection, the media was changed. Virus supernatant was harvested 48h post-transfection, filtered through a 0.45µm PVDF filter, concentrated with Lenti-X Concentrator (Clontech), aliquoted, and stored at -80°C.

5×10^5^ CTCs were transduced with lentivirus in 6-well plates in 2mL media supplemented with 6µg/mL Polybrene. 24h after infection, the media was changed. 72h after infection, cells were selected using puromycin (2µg/mL) for 3 days. Lentiviral titers were determined by infected cells with 6 different volumes of lentivirus and counting the number of surviving cells after 7 days of selection.

### CRISPR inactivation screen – mice injection and organ harvest

Brx-142 cells stably expressing GFP, luciferase, and KRAB-dCas9 (Addgene #96918) were transduced with the human CRISPR Inhibition Pooled Library (Dolcetto pool A, Addgene #92385) (12), at a MOI of 0.3, and fully selected with puromycin (2µg/mL), to achieve a representation of 300-350 cells per sgRNA. At the same time, plasmid DNA used for viral transduction (pDNA) was isolated as an input baseline distribution of guides. Immediately following selection, 8 female NSG (NOD. CgPrkscid Il2rgtm1Wjl/SzJ) mice were injected with 3×10^6^ cells each, via the tail vein. Mice were anesthetized with isoflurane, and a 90-day release 0.72mg estrogen pellet (Innovative Research of America) was implanted subcutaneously behind the neck of each mouse. After three months, the mice were sacrificed, and the whole lungs were harvested. All mouse handling was completed in compliance with ethical regulations and approved by the IACUC animal protocol 2010N000006. Lungs were divided into 25µg chunks and homogenized using a TissueLyser II (Qiagen), and DNA was extracted using NucleoSpin Tissue DNA extraction columns (Macherey-Nagel). PCR was performed by the Broad Institute Screening Plateform, and products were sequenced on the Illumina MiSeq platform.

### CRSIPR inactivation Screen - Analysis

Guide counts from all 8 mice were pulled together and were normalized to the total counts. Guide distribution was compared to the plasmid input distribution of guides, resulting in a fold change value for each guide. Guides with a count input lower than 100 were excluded. The most enriched guide for each gene was determined and corresponding genes were rank-ordered based on their fold enrichment, with a rank of 1 denoting most enriched.

### Stable Knock-down of HECTD4 and COX-2, with lentiviral production and infection

For stable knockdown of *HECTD4* and *COX-2* expression, pLKO.1-Puro lentiviral constructs expressing shRNA against human *HECTD4* and *COX-2,* respectively, were obtained from the MGH Molecular profiling laboratory (MPL) at the MGH Cancer Center.

For lentiviral production of shRNA, HEK 293T cells were transfected with the specific shRNA lentiviral constructs, specified below, together with pMD2.G (Addgene#12259) and psPAX2 (Addgene#12260) packaging plasmids using Lipofectamine 2000 reagent (Invitrogen). 48-72 h after transfection, culture medium (containing lentiviral particles) was collected, filtered through a 0.45µm PVDF filter, aliquoted and stored at -80°C, as described above. Viral particles were added to the MDA-MB-231, HCC1143, BT-549 and MDA-MB-468 cells in presence of polybrene (Santa Cruz, 8 µg/ml as final concentration) overnight. Cells were then selected with puromycin (2µg/mL) for 72 hours or hygromycin (500µg/mL) for 6 days.

The *HECTD4* mRNA sequences targeted with lentiviral shRNA clones are listed in **table S1**.

### Transient Knock-down of MKK7 and COX-2, with siRNA

For transient KD of *COX-2* or *MKK7*, MDA-MB-231 in monolayer cultures were treated with a SMARTpool comprised of 4 different siRNAs targeting COX-2 (Horizon Discovery # L-004557-00-0005), MKK7 (Horizon Discovery # L-004016-00-0005) or control (non-targeting pool, Horizon Discovery # D-001810-10-05). The cells were seeded into 6-well plates (1.5 x105 cells/well), and treated with 0.625 μL of siRNA 20µM, mixed with 6uL of lipofectamine RNAiMAX (Thermo Scientific #13778075) in reduced serum medium (Opti-MEM, Thermo Scientific #31985062), according to manufacturer’s instructions. The cells were placed in suspension after 24 hours and/or analyzed at the indicated time point.

### Mouse homogenous experiment

GFP+/luciferase+ MDA-MB-231 were infected with either scrambled vector (Addgene #136035) or shHECTD4 #1 and fully selected with puromycin (2µg/mL). For the injections, 2×10^5^ cells of each group, respectively were further combined into 1:1 Growth Factor Reduced Matrigel (Corning). The cell media (100uL total volume) was injected into the right inferior mammary fat pad of 4 female NSG mice for the control and *HECTD4*-KD groups. At day 34, the mice were sacrificed, and the primary tumors were harvested. Tumor weight and volume of the NSG mice xenograft was measured at day 34.

### Mouse mixing experiment-Validating the HECTD4 KD

GFP+/luciferase+/ mCherry-MDA-MB-231 were infected with scrambled vector (Addgene #136035) (GFP-shControl; *HECTD4*-WT) and GFP+/luciferase+/ mCherry+ MDA-MB-231 were infected with shHECTD4 (sh #1) (GFP-mCherry-*HECTD4*-KD) or with the scrambled vector (GFP-mCherry-shControl; *HECTD4*-WT). Cells were fully selected with puromycin (2µg/mL). GFP-shControl cells were mixed at ratio 1:1 with the GFP-mCherry-shControl cells , and a second mixing sample combined GFP-shControl cells and GFP-mCherry-*HECTD4*-KD cells. An aliquot of both mixes was saved for FACS analysis of the initial injection ratio. For the injections, 2×10^5^ cells of the mixes were further combined into 1:1 Growth Factor Reduced Matrigel (Corning #354230). The cell media (100uL total volume) was injected into the right inferior mammary fat pad of 4 female NSG mice for the control mix and 5 for the *HECTD4*-KD mix. Metastatic growth was measured weekly via in vivo imaging using the IVIS Lumina II (PerkinElmer) following intraperitoneal injection of D-luciferin (Sigma). We performed a survival surgery at day 36 to harvest the primary tumors. At day 62-65, mice were sacrificed, and lungs and livers were harvested. Fluorescent pictures of the organs were taken with an MVX10 Macro Zoom Fluorescence microscope (Olympus) with

GFP and mCherry filter. One fraction was stored into 10% formalin for 24h for fixation prior to immunohistochemistry. The rest of the organ was mechanically dissociated, followed by a biological digestion with collagenase/hyaluronidase in DMEM (StemCell Technologies Inc. # 07912) by rotation for 3 hours at 37°C. Digested lysates was cleared by centrifugation and rinsed in PBS. One liver sample in the *HECTD4*-KD group was excluded from the final analysis due to tissue and cellular damage as a result of the digestion process. In total, n = 4 for WT and n = 5 for *HECTD4*-KD for primary tumors; n = 4 for WT and n = 5 for *HECTD4*-KD for the lungs; n = 4 for WT and n = 4 for *HECTD4*-KD for the livers. Single cells were filtered through a mesh strainer and analyzed using an BD LSRFortessa X-20 flow cytometer (BD Bioscience) and a SH800S (Sony sorter). Cytometry data were processed with FlowJo software (version 10.8.1).

### Mouse mixing experiment-Double KD (HECTD4 and COX-2)

GFP +/luciferase+ MDA-MB-231 cells were infected with the mCherry (Addgene #176016) and the scrambled vector (Addgene #136035) as control (GFP-mCherry-shControl; *HECTD4*-WT and *COX-2*-WT). This control group was mixed with the following four groups:

1. GFP+/luciferase+/mCherry - MDA-MB-231 cells infected with scrambled vector (Addgene #136035) (GFP-shControl; *HECTD4*-WT and *COX-2* WT) ;
2. GFP+/luciferase+/mCherry - MDA-MB-231 cells infected with shCOX-2 (sh #1 and sh #2 mixed in a 1:1 ratio) (GFP-*COX-2*-KD; *HECTD4*-WT and *COX-2* KD);
3. GFP+/luciferase+/mCherry - MDA-MB-231 cells infected with shHECTD4 (sh #1, sh #2 and sh #3 mixed with a 1:1:1 ratio) ) (GFP-*HECTD4*-KD; *HECTD4*-KD and *COX-2* WT); and
4. GFP+/luciferase+/mCherry - MDA-MB-231 cells infected with shHECTD4 (sh #1, sh #2 and sh #3 mixed with a 1:1:1 ratio) and with shCOX-2 (sh #1 and sh #2 mixed in a 1:1 ratio) (GFP- Double-KD; *HECTD4*-KD and *COX-2* KD).

Cells were fully selected with puromycin (2µg/mL). GFP-mCherry-shControl cells were mixed with i. (GFP-shControl), ii. (GFP-*COX-2*-KD), iii. (GFP-*HECTD4*-KD) and iv. (GFP-Double-KD) at a ratio of 1:1. An aliquot of all four mixes was saved for FACS analysis of the initial injection ratio. For the injections, 2×10^5^ cells of the mixes were further combined into 1:1 Growth Factor Reduced Matrigel (Corning #354230). The cell:Matrigel mixes (100uL total volume) were injected into both the right and left inferior mammary fat pad of female NSG mice (n=6 mice per group; total of 24 mice). Primary tumor and metastatic growth were measured weekly via in vivo imaging using the IVIS Lumina II (PerkinElmer) and Aura Image Analysis Software by Spectral Instruments Imaging (version 4.0.8) following intraperitoneal injection of D-luciferin (Sigma #L6153). We performed survival surgeries on four out of the six mice in each group at days 24-25 to harvest the primary tumors. Two mice from each group were sacrificed and their primary tumors as well as lungs and livers were collected to assess for early metastasis development in the these organs. The volumes of the primary tumors were measured at this time. At day 54-55, the remaining four mice per group were sacrificed, and their lungs and livers were harvested. One fraction was stored into 10% formalin for 24h for fixation prior to immunohistochemistry. The rest of the organ was mechanically dissociated, followed by a biological digestion with collagenase/hyaluronidase in DMEM (StemCell Technologies Inc. # 07912) by rotation for 3 hours at 37°C. Digested lysates was cleared by centrifugation and rinsed in PBS. The 15 out of the 16 primary tumors of the eight mice sacrificed during days 24-25, were included in the final analysis (two mice per group, one primary tumor sample was excluded from the *COX-2* KD group due to tissue and cellular damage during digestion), however, their lungs and livers were excluded due to the absence of metastatic cells. Of the remaining 16 mice (4 mice per group), one primary tumor sample in the Double KD group, one lung sample in the *HECTD4* KD group, one lung sample in the Double KD group and one liver sample in the WT group were excluded from the final analysis due to tissue and cellular damage as a result of the digestion process. In total, n = 12 for WT, n = 11 for *COX-2*-KD, n = 12 for *HECTD4*-KD and n = 12 for Double KD for primary tumors; n = 4 for WT, n = 4 for *COX-2*-KD, n = 3 for *HECTD4*-KD and n = 3 for Double KD for the lungs; n = 3 for WT, n = 4 for *COX-2*-KD, n = 4 for *HECTD4*-KD and n = 4 for Double KD for the livers. Single cells were filtered through a mesh strainer and analyzed using a SH800S (Sony sorter). Cytometry data were processed with FlowJo software (version 10.8.1).

### Proliferation assays

For metabolic activity assays, CellTiter Glo reagent (Promega) was used according to the manufacturer’s instructions. Cells were seeded at a density of 3,000 cells per well in 3 or 6 replicates into 96-well Flat Bottom TC-treated Microplate (for adherent conditions) or into 96-well clear ultra-low attachment microplate (for anchorage-independent suspension conditions) (Corning) and incubated for the mentioned time prior to reading. Luminescence was measured with a Spectra Max M5 plate reader (Molecular Devices). All reads were normalized first to their day of plating and to the control of the plate.

### Colony Formation Assay in Soft agar

A mixture of 10,000 cells in assay medium and 0.3% agarose was seeded on to a solidified bed of 0.6% agarose on six-well plates. The plates were allowed to solidify at 4°C and incubated at 37°C. The cultures were fed once a week with 1mL of standard growth medium. Cells were incubated for 27 days and then stained with 500 μl of crystal violet stain (0.05% crystal violet, formaldehyde 1%, methanol 1%) for 30 minutes and washed with PBS. Colonies (>50 μm in diameter) were imaged using the MVX10 Macro Zoom Fluorescence microscope (Olympus) with a brightfield and counted with the HALO software.

### Prime-editing ALFA-tagging of endogenous HECTD4

MDA-MB-231 cells were seeded into 24-well flat-bottom cell culture plates (Corning) for PE treatment at 30,000 cells/well. Transfections were carried out 24 h post-seeding with 4.5 ug PE2 plasmid (Addgene plasmid (PE2) #132775) , 1.5 ug pegRNA, and 500 ng ngRNA plasmid per transfection (per well, in a 24-well plate). TransIT-X2 (Mirus, MIR #6004) was used as the lipofection reagent at 0.3 mL per transfection. The sequences used are listed in **table S2.**

### RNA extraction, cDNA synthesis and quantitative real-time PCR

RNA extracted from cultured breast cancer cells was prepared using the RNeasy Mini kit (QIAGEN) with DNase I digestion on the column) according to the manufacturer’s instruction. cDNA was synthesized from 100-500 ng RNA using SuperScript IV VILO Master Mix (Invitrogen # 11756050). qPCR was performed using the primers listed in **table S3**. qRT-PCR was run by QuantumStudio 5 instrument (ThermoFisher Scientific). Each reaction was performed in 3 or 6 replicates and fold expression change was calculated using the comparative DDCt method, normalized to PP1A or GAPDH endogenous controls.

### Protein cell lysate Preparation and Western blot

Cells were washed in ice-cold PBS and lysed in RIPA buffer (Thermo Scientific #NC9484499) or Triton X-100 lysis buffer (Thermo Scientific #J62289.AK) containing a cocktail protease inhibitor (Halt^TM^ Protease Inhibitor Cocktail, Thermo Scientific #78429). The lysate was incubated on ice for 20 minutes and cleared by centrifugation at 4 °C for 10 minutes at 13’000 RPM. The supernatant was used for protein sample preparation. Protein concentration was determined using DC protein assay (Bio-Rad, # 5000122). Samples were prepared diluted in PBS together with loading and reducing agents and heated at 95°C for 5 min. Proteins (between 10ug and 20 ug) were separated on 4-15% polyacrylamide gradient-SDS gels (Bio-Rad # 5671084), on 4-20% (Bio-Rad #4561093) and 7.5% (Bio-Rad # 4561023) and transferred onto nitrocellulose membranes using the iBlot 2 Gel Transfer Device (ThermoFisher). After blocking with PBS-T with 5% milk for 1 hour at room temperature, membranes were incubated with primary antibodies overnight at 4 °C at the recommended concentrations. The primary antibodies are listed in the **table S4.** After 3 washes of 5 minutes in PBS-T, HRP conjugated secondary antibodies (1:10,000; Bio-Rad; Cat#5196–2504) were applied at room temperature. Blots were washed in PBS-T 3 x 5 min and incubated with the enhanced chemiluminescence substrate (Bio-Rad # 1705062), before being digitally imaged on a chemiluminescent ChemiDoc Imaging System MP (Bio-Rad #1705061).

### Proteomics: Cell lysis, protein digestion, and TMT labeling

*HECTD4*-KD (shHECTD4 #3, n=3) and scrambled control MDA-MB-231 (n=3) cells were seeded in ultra-low adherent plates (Corning) for four days. The proteasome inhibitor MG132 (Selleck #S2619, final concentration of 20 µM) and lysosomal inhibitor (Bafilomycin A1: Sigma Aldrich #B1793, final concentration of 20 nM) were added to the cells, at the indicated concentrations, for 6 hours. Cell pellets were processed as previously described (38, 39). Briefly, cell pellets were lysed with 1000 µL of lysis buffer (75 mM NaCl, 3 % SDS, 1 mM NaF, 1 mM beta-glycerophosphate, 1 mM sodium orthovanadate, 10 mM sodium pyrophosphate, 1 mM PMSF and 1X Roche Complete Mini EDTA free protease inhibitors in 50 mM EPPS, pH 8.5). Proteins were then reduced with 5 mM DTT at 56°C for 30 min and alkylated with 15 mM iodoacetamide in the dark at room temperature for 20 min. Tricholoracetic acid (TCA) was used to precipitate the reduced and alkylated proteins. Proteins were solubilized in 1M urea in 50mM EPPS, ph 8.5 and digested in a two-step process (overnight digestion at room temperature with 1 µg/µL of Lys-C (Wako) followed by six hours of digestion with trypsin (sequencing grade, Promega) at a final concentration of 1 ng/μL for 6 hr at 37 °C). The digest was acidified with 10 % TFA and peptides were desalted on C18 solid-phase extraction (SPE) (Sep-Pak, Waters) columns. The concentration of the desalted peptide solutions was measured with a BCA assay, and peptides were aliquoted keeping 1950ug for the diGly proteomics and 100ug for total proteome. The samples were dried under vacuum and stored at -80 °C until they were labeled with TMT reagents. For total proteomics, TMT labeling was performed with 18-plex tandem mass tag (TMT) reagents (Thermo Scientific) as described below (38). The TMT labeled peptides were pooled and desalted via C18 SPE and then fractionated into 24 fractions using Basic pH Reversed-Phase Liquid Chromatography (bRPLC). The individual fractions were analyzed in 3-hour runs via LC-MS2/MS3 on an Orbitrap FusionLumos mass spectrometer using the Simultaneous Precursor Selection (SPS) supported MS3 method (40, 41) .

For the diGly proteomics, the co-immunoprecipitation of KεGG-modified peptides was carried out according to manufacturer’s instruction (Cell Signaling PTMScan, #59322). Briefly, the samples were resuspended in the provided binding buffer. 20uL of bead-slurry were used per condition, and the samples were incubated on an end-over-end rotator for 2 hours at 4°C. Washes were performed with the provided wash buffer, and 2 rounds of elution with IAP Elution Buffer (0.15% TFA) were performed. A second round of co-immunoprecipitation was subsequently performed with fresh bead-lysate. Eluted DiGly-peptides were combined and labeled with TMT18plex reagents (Thermo Fisher Scientific). For labeling with TMT 18 plex reagents, 25 µg of peptides were dried and resuspended in 25 µL of 200 mM EPPS (pH 8.5), 30% acetonitrile (ACN). Labeling was performed by adding 70 μg TMT reagent in anhydrous ACN and incubating at room temperature for 1 h. The reaction was stopped by addition of 5% (w/v) hydroxylamine in 200 mM EPPS (pH 8.5) to a final concentration of 0.5% hydroxylamine and incubation at room temperature for 15 min. Samples were acidified with 1% TFA, and combined. The pooled samples were desalted using Sep-Pak C18 SPE cartridges and vacuum dried. The peptides were then fractionated using Pierce™ High pH Reversed-Phase Peptide Fractionation Kit (Thermo Scientific Cat 84868) according to the manufacturer’s instructions and vaccum dried. The dried fractions were resuspended in 5 % ACN/5 % formic acid and analysed with high resolution LC-MS2 on an Orbitrap FusionLumos mass spectrometer. MS2 spectra were assigned using a COMET-based in-house built proteomics analysis platform (42) allowing methionine oxidation (+15.99492 Da) and lysine ubiquitylation (+ 114.04293 Da) as a variable modification. A target-decoy database-based search was used to filter the false-discovery rate (FDR) of protein identifications of <1% (43). Peptides that matched to more than one protein were assigned to that protein containing the largest number of matched redundant peptide sequences following the law of parsimony. TMT reporter ion intensities were extracted from the MS3 spectra, selecting the most intense ion within a 0.003-*m*/*z* window centered at the predicted *m*/*z* value for each reporter ion, and spectra were used for quantification if the sum of the S/N values of all reporter ions divided by the number of analyzed channels was ≥10 and the isolation specificity for the precursor ion was ≥0.75. Protein intensities were calculated by summing the TMT reporter ions for all peptides assigned to a protein.

The full proteome data was normalized in a two-step process, a row (protein) normalization followed by a column (sample) normalization. The row normalization involved scaling the row intensity values by the global median of all row means divided by each rows mean, adjusting each row mean to equal the global median. Similarly, the column normalization involved scaling the column intensities so that each column median would be equal the average of all sample medians. The raw diGly peptide data was normalized in an analogous procedure.

### Proteomic Analysis

Pairwise Pearson correlations between replicate proteome profiles were generated and two samples, 7_Control_MG132_BafA1 (average Pearson= -0.06) and 12_KD_sh3_MG132_BafA1 (average Pearson=0.35) were removed from downstream analyses because the average correlation with their replicates was below 0.5. Differential analysis of total proteome expression levels between *HECTD4* knockdown and control sample was performed using the Limma R package (44) which uses a moderated t-statistic to generate a significance p-value for each protein. P-values were adjusted for multiple hypothesis testing by the Benjamini-Hochberg correction (45). Protein with at least 1.5 fold change and a corrected p-value < 0.25 were kept for further analysis.

Differential expression analysis for the triplicate diGly data was performed as described previously for the total proteome data. DiGly peptide levels were normalized to their full protein expression levels prior to differential expression analysis. Out of the total 3447 diGly peptides quantified, normalized expression levels for 3311 peptides were generated and used in downstream analyses.

### HECTD4 Short Active and Short Mutant assay

For the cloning of short active and mutant form of HECTD4, a DNA sequence coding for the aminoacids 3988 to 4428 of the *HECTD4* gene (C-terminus; short active with the regular sequence; the short mutant with the residue C4396 mutated into Alanine) in frame with the ALFA tag at its 5’ end and flanked by *BamHI* and *XbaI* inside the subcloning vector pUC-GW-Amp was ordered from GENEWIZ.

This fragment was subcloned by restriction (*BamHI* and *XbaI*) and ligation into the lenti-vector expression backbone plasmid pLenti-CMV-Blast-empty-(w263-1) (Addgene #125133). Sequence integrity was validated with Sanger sequencing.

Lenti-vectors were prepared in 293T cells as described above. Briefly, 293T cells were transfected with the lentiviral constructs, specified above, together with pMD2.G (Addgene#12259) and psPAX2 (Addgene#12260) packaging plasmids using Lipofectamine 2000 reagent (Invitrogen). 48-72 h after transfection, culture medium (containing lentiviral particles) was collected, filtered through a 0.45µm PVDF filter, aliquoted and stored at -80°C. MDA-MB-231 transduction was carried through an overnight incubation of lenti-vector supernatants with 8 µg/ml Polybrene (Santa Cruz). Transduced cells were selected for with 10 µg/ml Blasticidin S HCL (Gibco).

### In vitro transcription and translation

All *in vitro* synthesized substrates (COX-2 and MKK7) were cloned under the SP6 promoter in the pCS2+N-3HA cloning vector (Addgene #196653). The corresponding created plasmids are the pCS2-3HA-COX-2 and the pCS2-3HA-MKK7. The substrate was generated by incubating 3 μl (400 ng) of plasmid DNA in 25 μl of rabbit reticulocyte lysate (Promega, L2080) for 1 h at 30 °C. Reactions were terminated by rapid dilution with 1× PBS. Substrates were used for in vitro ubiquitylation assays.

### HECTD4 Immunoprecipitation/ purification

Cells were lysed in lysis buffer (TRIS pH 7.5 20µM, 1mM EDTA, 1mM EGTA, 150mM NaCl, 1% Triton X-100, protease and phosphatase inhibitors (ThermoFisher Cat#78440). Lysates were kept on ice for 20 minutes, and then cleared by centrifugation at 13,000 rpm for 15 minutes at 4°C. HECTD4 immunoprecipitation was performed using 500 μg to 1 mg of cellular lysates using magnetic beads crossed-linked to ALFA nanobody (Nanotag Biotechnologies, cat## N1515-L), previously rinsed in the aforementioned lysis buffer (40uL/mg of cellular lysis). 5% was stored as input sample.

Magnetic beads and lysis were incubated with rotation at 4°C for 2 hours. Beads were washed with ice cold lysis buffer four times. For in vitro ubiquitylation assay, magnetic bead-attached HECTD4 was used as it (5uL of beads per reaction). For co-immunoprecipitation analysis, samples were eluted with urea sample buffer (120 mM Tris 6.8; 4% SDS; 4M urea; 20% glycerol, 10mM βME). Elutes were then run and analyzed by western blots.

### In vitro Ubiquitination

In vitro ubiquitination assays were performed in a 15 μl reaction volume: 0.25 μl of 10 μM E1 (Fisher Scientific # E304050), 1 μl of 10 μM E2 (UBE2D) (Boston Biochem # E2-622), 1 μl of 10 mg/ml ubiquitin (Boston Biochem, U-100H), 1 μl of 15 mM DTT, 1.5 μl of energy mix (150 mM creatine phosphate, 20 mM ATP, 20 mM MgCl2 , 2 mM EGTA, pH to 7.5 with KOH), 4.25 μl of 1 × PBS, 1 μl of 10 × ubiquitylation assay buffer (250 mM Tris 7.5, 500 mM NaCl, and 100 mM MgCl2 ) and 1 μl of substrate (in vitro translated) were premixed and added to 4 μl of HECTD4-purified or empty magnetic beads (see ‘HECTD4 immunoprecipitation/purification’). Reactions were performed at 30 °C with shaking for 60 min, unless noted otherwise. For the deubiquitination reaction, 0.25 µL of the de-ubiquitinating enzyme Ubiquitin Specific Peptidase 2 (USP2) (Fisher Scientific # E504050) was added to the sample following incubation for 60 min at 30 °C. The samples with USP2 were incubated an additional 30 min at 30 °C. All reactions were stopped by adding 2× urea sample buffer and resolved on SDS-acrylamide gels.

### HECTD4 evolutionary tree and clinical data visualization

The evolutionary tree showing the 13 members of the HECT subfamily was developed using the Neighbor-Joining algorithm implemented in the MEGA-11 program using Poisson correction method (46). HECTD4 gene expression data (3691 normal samples, 29,376 tumors and 453 metastatic) was derived from gene chip-based studies of the Gene Expression Omnibus of the National Center for Biotechnology Information (NCBI-GEO), plotted on TNMplot.com (15). Proteomic data was obtained from the proteomic datasets of the National Cancer Institute’s Clinical Proteomic Tumor Analysis Consortium (CPTAC) and National Cancer Institute’s International Cancer Proteogenome Consortium (ICPC). Data was visualized by the interactive web server Cancer Proteogenomic Data Analysis Site (CProSite) (47). Kaplan Meier plot for progression-free survival was derived from the transcriptomic datasets with follow-up and clinical data from the GEO repository and were plotted with the interactive web server KMplot (16). HECTD4 gene expression levels in different breast cancer subtypes, the frequency of different HECTD4 mutations in breast cancer subtypes and HECTD4 mRNA expression in different tumor types were obtained from the pan-cancer data from The Cancer Genome Atlas (TCGA) (48). We used the pre-processed RNA sequencing gene expression file (EBPlusPlusAdjustPANCAN_IlluminaHiSeq_RNASeqV2.geneExp.tsv) from Pan-cancer atlas repository. The processed somatic mutation data was taken from the Pan-cancer atlas repository (mc3.v0.2.8.PUBLIC.maf.gz) which calculated by the MC3 collaboration aiming at a consensual and unified variant calling (49). We considered as somatically mutated if one of the following holds:

- The mutations were truncating (“Nonsense_Mutation”, “Frame_Shift_Del”, “Frame_Shift_Ins”, “Translation_Start_Site”, “Splice_Site”)
- The nonsynonymous mutations (“Missense_Mutation”, “In_Frame_Del”, “In_Frame_Ins”,“Nonstop_Mutation”) were “deleterious” in the SIFT field and “probably_damaging” in the PolyPhen.
- The nonsynonymous mutations were predicted to be “pathogenic” or “likely_pathogenic” by ClinVar (50).
- In any case an above mutation was labelled as “Benign” according to ClinVar we did not consider this mutation. Germline mutations were processed by other groups (51) and obtained from GDC data portal (PCA_pathVar_integrated_filtered_adjusted.tsv), and the variants of unknown significance were excluded. Copy number ASCAT calls are obtained through (https://github.com/VanLoo-lab/ascat/) (52) and loss-of-heterozygosity is defined as minor allele being 0.

### Correlation of HECTD4 and COX-2 across 21 Breast cancer cell lines

Publicly available cell line proteomics data was collected across 31 breast cell lines (19). A total of 21 breast cell lines were selected for which both the canonical HECTD4 and COX-2 proteins were quantified. A Pearson correlation test between HECTD4 and COX-2 across the cell lines was calculated using the cor.test function in R (53).

### Statistical analysis

GraphPad Prism software (version 10.2.2) was used to generate graphs and to perform the corresponding statistical tests indicated in the figure legends.

### Illustrations

Illustrations were created with BioRender.com.

### Data Availability

All mass spectrometry RAW data can be accessed through the MassIVE data repository (massive.ucsd.edu) under the accession number MSV000096318. Data can be accessed using the accession number “MSV000096318_reviewer” and the password “Joanna_2024”.

## Author Contributions

J.A.V., C.T., E.O., S.M. and D.A.H. conceived the project, provided leadership for the project, and drafted the manuscript. J.A.V., C.T., D.S.M., R.Y.E., S.A., R.M., S.H., Z.J.N., H.C.R., E.F.Z., J.G., R.N.S., and E.A. conducted all the experiments. J.A.V., C.T., D.S.M., R.Y.E., S.A., R.M., S.H., B.S.W., B.K.W., E.A., D.B.F., M.Y. and D.C.G. analyzed all the data. J.A.V., C.T., D.C.G., A.E.H.E., W.H., E.O., S.M. and D.A.H. interpreted the results. All authors reviewed and edited the manuscript.

## Acknowledgments

We thank the patients and their families and the medical teams involved in caring for the patients. We thank L. Libby for technical support. We thank all lab members in D.A.H./S.M.’s lab for discussions. This work was supported by grants from National Institute of Health (2RO1CA129933 to D.A.H.; U01EB012493 to D.A.H., and S.M.; R35GM118158 to J.K.J.), Howard Hughes Medical Institute (to D.A.H.), ESSCO Breast Cancer Research Fund (to S.M.), Breast Cancer Research Foundation (to D.A.H.), National Foundation for Cancer Research (to D.A.H.) and the Placide Nicod Foundation (to J.A.V.). The content is solely the responsibility of the authors and does not necessarily represent the official views of the NIH. These funders had no role in the study design, data collection and analysis, decision to publish or preparation of the manuscript.

