## Supplemental Information for "The E3 ligase HECTD4 regulates COX-2 dependent tumor progression and metastasis"

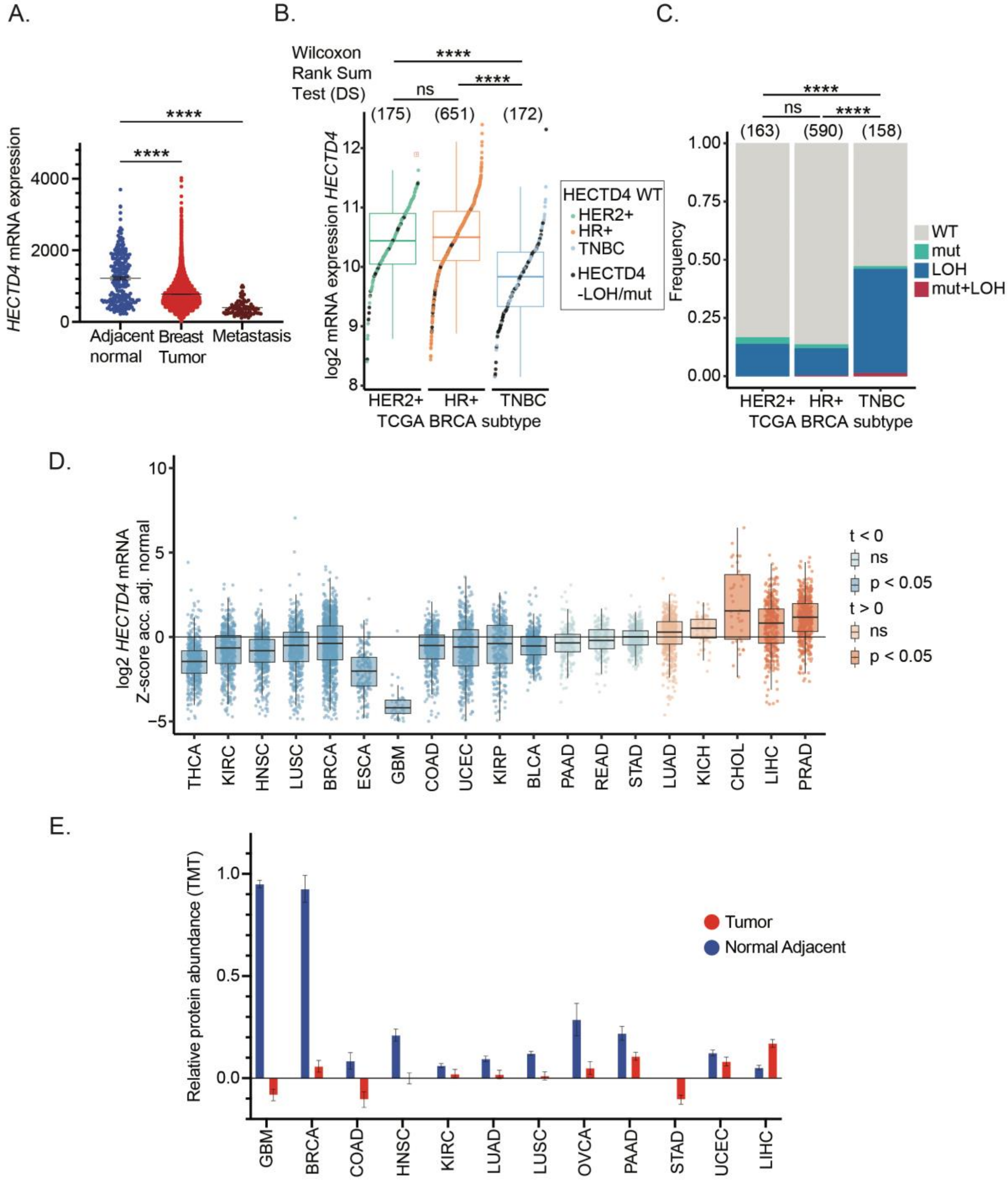

A.
B.
C.
D.
E.
HECTD4 mRNA expression
4000
2000
0
****
Adjacent normal
Breast Tumor
Metastasis
Wilcoxon Rank Sum Test (DS)
ns
(175) (651) (172)
log2 mRNA expression HECTD4
HECTD4 WT
HER2+
HR+
TNBC
HECTD4 -LOH/mut
HER2+ HR+ TNBC
TCGA BRCA subtype
(163) (590) (158)
Frequency
WT
mut
LOH
mut+LOH
log2 HECTD4 mRNA Z-score acc. adj. normal
t < 0
ns
p < 0.05
t > 0
THCA KIRC HNSC LUSC BRCA ESCA GBM COAD UCEC KIRP BLCA PAAD READ STAD LUAD KICH CHOL LIHC PRAD
Relative protein abundance (TMT)
Tumor
Normal Adjacent
GBM BRCA COAD HNSC KIRC LUAD LUSC OVCA PAAD STAD UCEC LIHC

**Fig. S1. HECTD4 expression in clinical datasets (A)** *HECTD4* gene expression is significantly decreased in primary and metastatic tumors compared with normal breast tissue (3691 normal breast tissues, 29,376 primary breast cancers and 453 metastatic breast cancers). Significance was calculated with one-way ANOVA test, with Dunnett's multiple comparison test. **(B)** *HECTD4* mRNA expression levels are compared across breast cancer subtypes with Wilcoxon Rank Sum test and p-values shown on top; loss of heterozygosity (LOH) or damaging/pathogenic point mutations are marked in black. **(C)** Frequency of *HECTD4* LOH and truncating, damaging or pathogenic single nucleotide variants (SNVs) and indels (SIFT or PolyPhen damaging, or ClinVar pathogenic or likely pathogenic) are shown in each breast cancer subtype. Fisher t-test was used to compare samples that are wild type (WT) or has mutation or LOH, p-values are shown on top of the bar plot. **(D)** *HECTD4* mRNA expression in each tumor type was Z-score normalized according to adjacent normal tissue, where negative value indicates a downregulation in tumor and a higher value indicates an upregulation; Z-score normalization is performed by subtracting the mean and dividing by the standard deviation in adjacent normal samples (tumor types with >3 adjacent normal samples are shown). T-test was performed, and p-values indicated by the color of boxes and markers were adjusted with the Benjamini-Hochberg procedure. Tumor types are sorted according to p-value, and the sign of the t-value with those with higher significance are at the two extremities. THCA: Thyroid cancer, KIRC: Kidney renal clear cell carcinoma, HNSC: Head and neck squamous cell carcinoma, LUSC: Lung squamous cell carcinoma, BRCA: Breast cancer, ESCA: Esophageal cancer, GBM: Glioblastoma, COAD: Colon adenocarcinoma, UCEC: Uterine corpus endometrial carcinoma, KIRP: Kidney renal papillary cell carcinoma, BLCA: Bladder urothelial carcinoma, PAAD: Pancreatic adenocarcinoma, READ: Rectum adenocarcinoma, STAD: Stomach adenocarcinoma, LUAD: Lung adenocarcinoma, KICH: Chromophobe renal cell carcinoma, CHOL: Cholangiocarcinoma, LIHC: Liver hepatocellular carcinoma, PRAD: Prostate adenocarcinoma **(E)** HECTD4 protein levels across different normal tissues and tumor types. Error bars represent mean +/- SD. GBM: Glioblastoma, BRCA: Breast cancer, COAD: Colon adenocarcinoma, HNSC: Head and neck squamous cell carcinoma, KIRC: Kidney renal clear cell carcinoma, LUAD: Lung adenocarcinoma, LUSC: Lung squamous cell carcinoma, OVCA: Ovarian carcinoma, PAAD: Pancreatic adenocarcinoma, STAD: Stomach adenocarcinoma, UCEC: Uterine corpus endometrial carcinoma, LIHC: Liver hepatocellular carcinoma. (* $p<0.05$, ** $p<0.01$, *** $p<0.001$, **** $p<0.0001$)

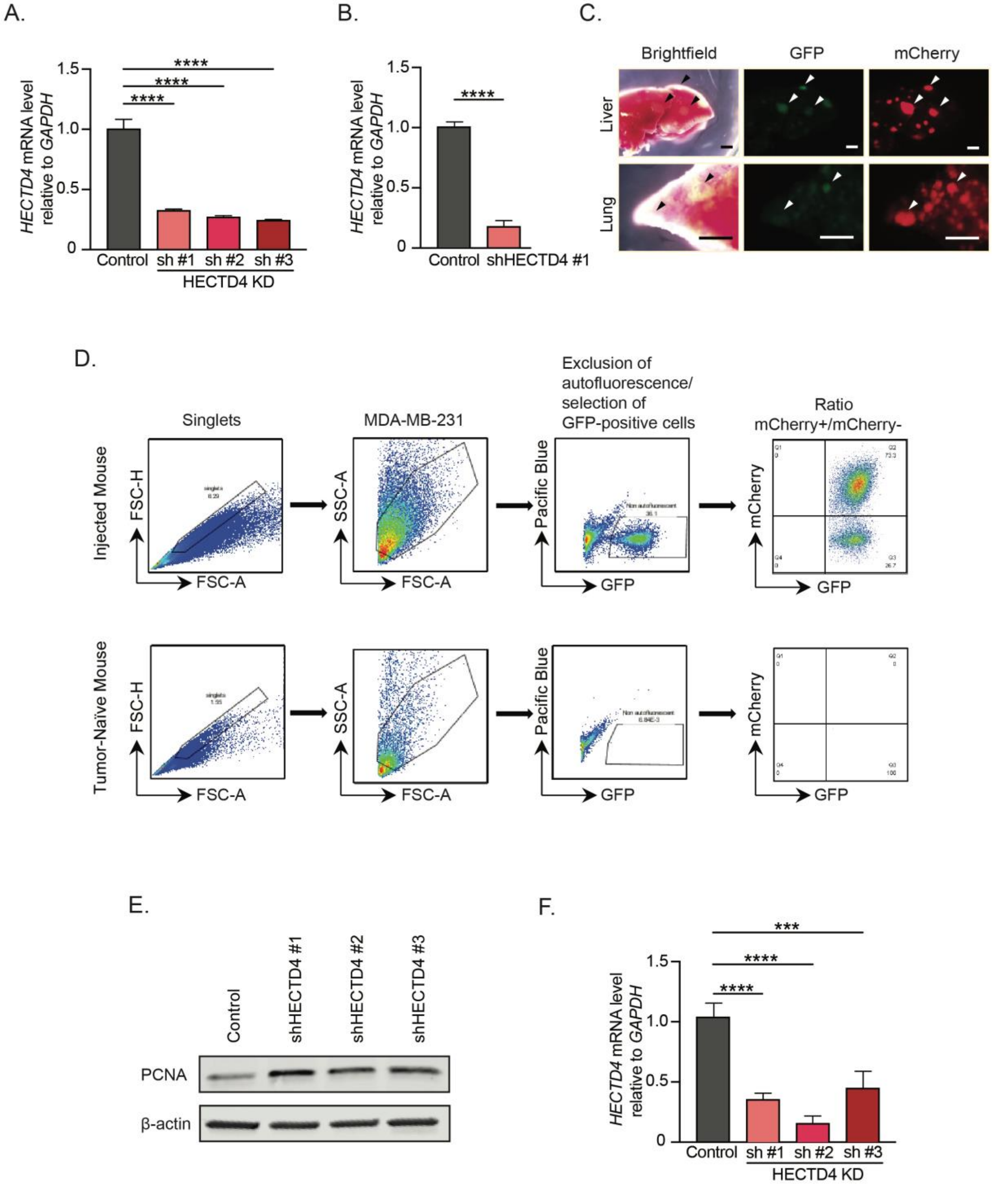

A.
HECTD4 mRNA level relative to GAPDH
1.5
1.0
0.5
0
****
****
****
Control
sh #1
sh #2
sh #3
HECTD4 KD
B.
HECTD4 mRNA level relative to GAPDH
****
Control
shHECTD4 #1
C.
Brightfield
GFP
mCherry
Liver
Lung
D.
Singlets
MDA-MB-231
Exclusion of autofluorescence/ selection of GFP-positive cells
Ratio mCherry+/mCherry-
Injected Mouse
Tumor-Naive Mouse
FSC-H
FSC-A
SSC-A
Pacific Blue
GFP
mCherry
E.
Control
shHECTD4 #1
shHECTD4 #2
shHECTD4 #3
PCNA
β-actin
F.
HECTD4 mRNA level relative to GAPDH
***
****
****
Control
sh #1
sh #2
sh #3
HECTD4 KD

**Fig. S2. HECTD4 depletion increases tumor cell proliferation *in vivo* and under anchorage-independent conditions (A)** shRNA-mediated *HECTD4* knockdown in MDA-MB-231 leads to reduced *HECTD4* mRNA levels compared to control cells transduced with scramble shRNA. Cells infected with shHECTD4 #1 were used for the experimental result shown **Fig. 2A.** Cells in which HECTD4 was knocked down with all three viruses were used for experiments shown in **Fig. 2C.** Error bars represent mean +/- SD. Significance was calculated using one-way ANOVA test, with Dunnett's multiple comparison test. **(B)** shRNA-mediated *HECTD4* knockdown in MDA-MB-231 leads to reduced *HECTD4* mRNA levels compared to control cells transduced with scramble shRNA. These cells were used for the *in vivo* mixing experiment in **Fig. 2B.** Error bars represent mean +/- SD. Significance was calculated using unpaired student t-test. **(C)** Representative images of the liver and lungs from the experiment shown in **Fig. 2B** with macro-metastasis, in brightfield and the specific emission filter to expose GFP-fluorescent cells and mCherry-fluorescent cells. Tumor cells are indicated by arrow heads. Scale bar: 2 mm. **(D)** Flow cytometry gating procedure for the detection of GFP+/mCherry+ tumor cells in the tumor cell mixtures in the experiment indicated in **Fig. 2B**. The gating strategy is specific, with no tumor cells detected in a naïve mouse (lower line). **(E)** *HECTD4*-KD cells compared with control cells have increased levels of PCNA protein in suspension. Cells were cultured in suspension for 3 days before the protein levels were analyzed by Western blot. **(F)** shRNA-mediated *HECTD4* knockdown in MDA-MB-231 leads to reduced *HECTD4* mRNA levels compared to control cells transduced with scramble shRNA. These cells have been used in the soft agar experiments shown in **Fig. 2D.** Error bars represent mean +/- SD. Significance was calculated using one-way ANOVA test, with Dunnett's multiple comparison test. (* $p<0.05$, ** $p<0.01$, *** $p<0.001$, **** $p<0.0001$)

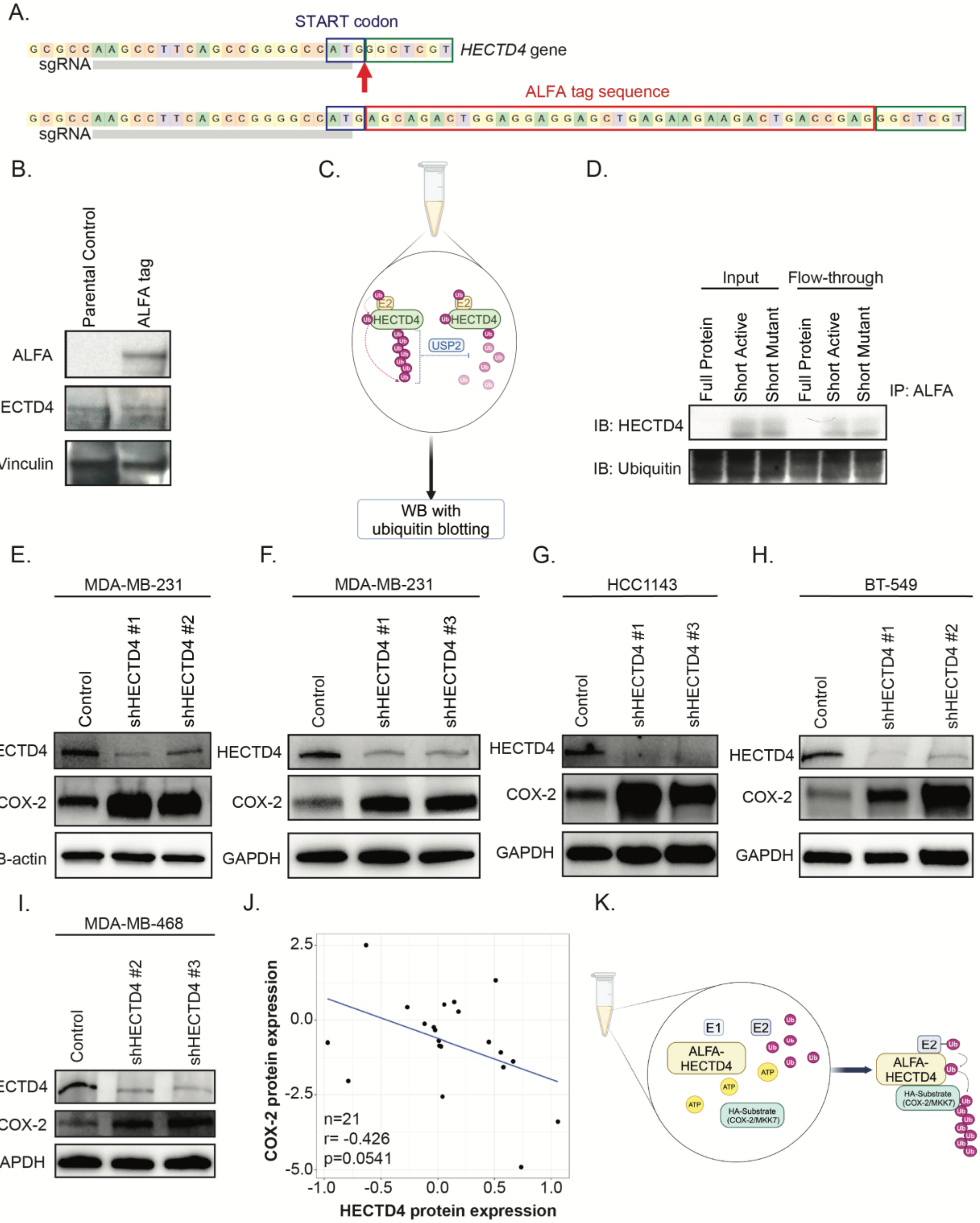
A.
START codon
G C G C C A A G C C T T C A G C C G G G G C C A T G G G C T C G T
HECTD4 gene
sgRNA
ALFA tag sequence
G C G C C A A G C C T T C A G C C G G G G C C A T G A G C A G A C T G G A G G A G G A G C T G A G A A G A A G A C T G A C C G A G G G C T C G T
sgRNA
B.
Parental Control
ALFA tag
ALFA
HECTD4
Vinculin
C.
E2
HECTD4
USP2
Ub
WB with ubiquitin blotting
D.
Input
Flow-through
Full Protein
Short Active
Short Mutant
Full Protein
Short Active
Short Mutant
IP: ALFA
IB: HECTD4
IB: Ubiquitin
E.
MDA-MB-231
Control
shHECTD4 #1
shHECTD4 #2
HECTD4
COX-2
B-actin
F.
MDA-MB-231
Control
shHECTD4 #1
shHECTD4 #3
HECTD4
COX-2
GAPDH
G.
HCC1143
Control
shHECTD4 #1
shHECTD4 #3
HECTD4
COX-2
GAPDH
H.
BT-549
Control
shHECTD4 #1
shHECTD4 #2
HECTD4
COX-2
GAPDH
I.
MDA-MB-468
Control
shHECTD4 #2
shHECTD4 #3
HECTD4
COX-2
GAPDH
J.
COX-2 protein expression
HECTD4 protein expression
2.5
0.0
-2.5
-5.0
-1.0
-0.5
0.0
0.5
1.0
n=21
r= -0.426
p=0.0541
K.
E1
E2
ALFA-HECTD4
ATP
HA-Substrate (COX-2/MKK7)
Ub

**Fig. S3. Ubiquitination and degradation of COX-2 by HECTD4 (A)** The endogenous tagging of HECTD4 at the N-terminus with prime-editing CRISPR strategy (1). The plot shows the integration of the exogenous 27-bp ALFA sequence at the 3' of the START codon of the HECTD4 sequence. **(B)** Western blotting against the ALFA tag, detects HECTD4. Parental cells are shown as control. Western blotting for vinculin is shown as loading control. **(C)** Schematic representation of HECTD4-dependent ubiquitin chain formation in vitro. **(D)** The lysates from cells transfected with the short-active and short-mutant forms of alfa-HECTD4 were flowed through an ALFA-tag activated column. The input lysate, flow- through and the eluate were blotted and probed with antibodies against HECTD4 and ubiquitin. Untransfected parental cells are shown as control. The input lysate and flow-through are shown here, the elute is shown in **Fig. 3B (Lower)**. **(E)** Western blot showing increased COX-2 protein in *HECTD4*-KD (sh #1 and sh #2) MDA-MB-231 cells growing in suspension. **(F)** Western blot showing increased COX-2 protein in *HECTD4*-KD (sh #1 and sh #3) MDA-MB-231 cells growing in suspension. **(G)** Western blot showing increased COX-2 protein in *HECTD4*-KD (sh #1 and sh #3) HCC1143 cells growing in suspension. **(H)** Western blot showing increased COX-2 protein in *HECTD4*-KD (sh #1 and sh #2) BT-549 cells growing in suspension. **(I)** Western blot showing increased COX-2 protein in *HECTD4*-KD (sh #2 and sh #3) MDA-MB-468 cells growing in suspension. **(J)** Analysis of publicly available proteomics data from 21 human breast cancer cell lines (2) shows a negative correlation between HECTD4 and COX-2 protein levels (N = 21, r = -0.426, p = 0.0541). A Pearson correlation test between HECTD4 and COX-2 across the cell lines was calculated. **(K)** Schematic representation of HECTD4-dependent ubiquitin chain formation of the HA- tagged proteins in vitro. HA-tagged COX-2 and MKK7 were produced by *in vitro* transcription-translation in reticulocytes and incubated with active HECTD4, immunoprecipitated from freshly lysed cells using ALFA-tagged beads. E1, E2, ATP, HA-tagged COX-2 or MKK7, ALFA-tagged HECTD4 and ubiquitin were added into the reaction. The ubiquitination of the HA-tagged protein of interest was detected by western blotting.

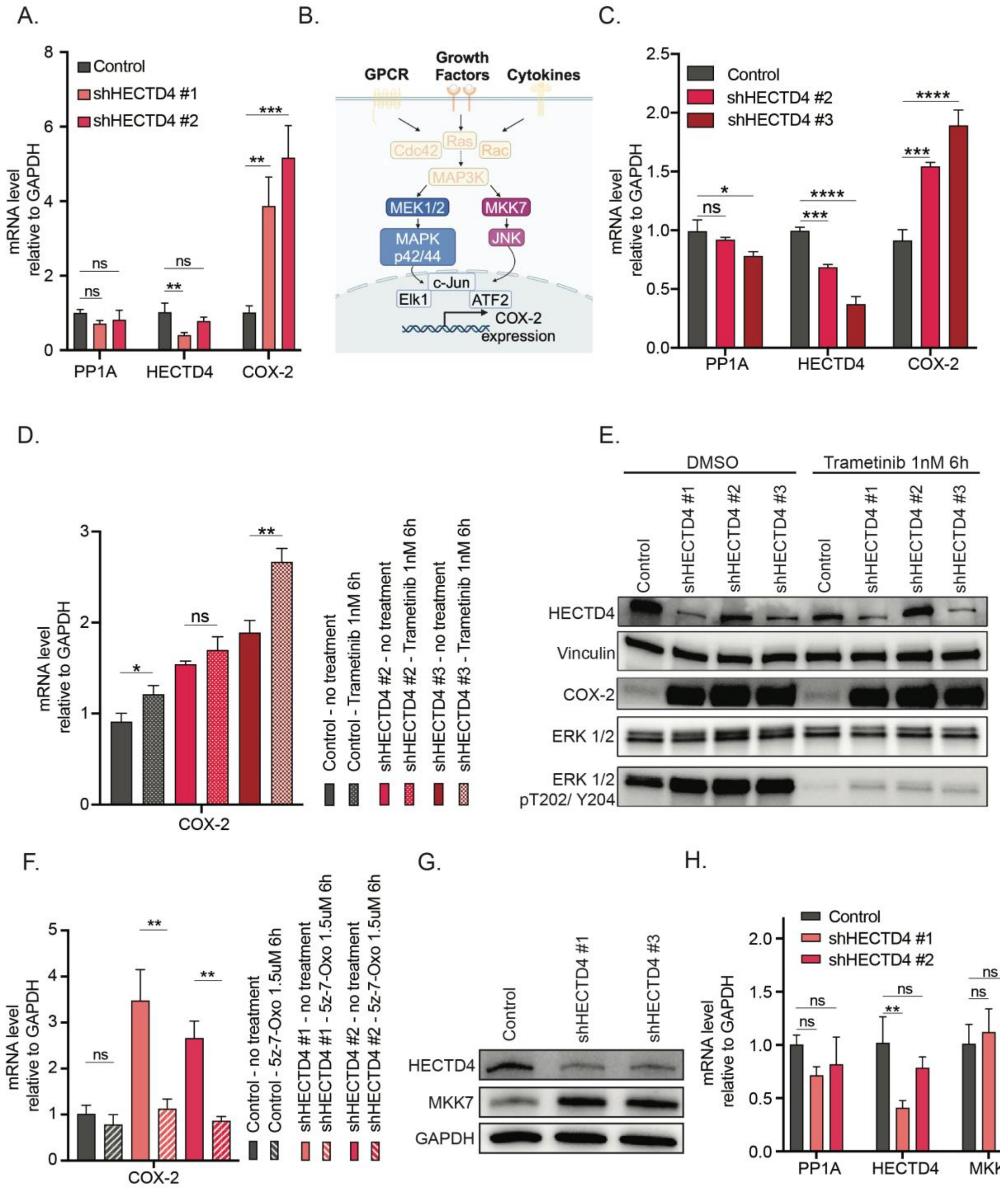
A.
Control
shHECTD4 #1
shHECTD4 #2
mRNA level relative to GAPDH
PP1A
HECTD4
COX-2
B.
GPCR
Growth Factors
Cytokines
Cdc42
Ras
Rac
MAP3K
MEK1/2
MKK7
MAPK p42/44
JNK
c-Jun
Elk1
ATF2
COX-2 expression
C.
Control
shHECTD4 #2
shHECTD4 #3
D.
Control - no treatment
Control - Trametinib 1nM 6h
shHECTD4 #2 - no treatment
shHECTD4 #2 - Trametinib 1nM 6h
shHECTD4 #3 - no treatment
shHECTD4 #3 - Trametinib 1nM 6h
E.
DMSO
Trametinib 1nM 6h
Control
shHECTD4 #1
shHECTD4 #2
shHECTD4 #3
HECTD4
Vinculin
COX-2
ERK 1/2
ERK 1/2 pT202/ Y204
F.
Control - no treatment
Control - 5z-7-Oxo 1.5uM 6h
shHECTD4 #1 - no treatment
shHECTD4 #1 - 5z-7-Oxo 1.5uM 6h
shHECTD4 #2 - no treatment
shHECTD4 #2 - 5z-7-Oxo 1.5uM 6h
G.
MKK7
GAPDH
H.
MKK7

**Fig. S4. HECTD4 targets the COX-2 regulator MKK7 (A)** qRT-PCR of *PP1A*, *HECTD4* and *COX-2* mRNA in control and *HECTD4*-KD cells, showing that *COX-2* mRNA levels rise upon *HECTD4* depletion. Error bars represent mean +/- SD. Significance was calculated using one-way ANOVA test, with Dunnett's multiple comparison test. **(B)** Diagram showing the MAPK pathway regulating COX-2 transcription; selected players are highlighted. Several stimuli, including growth factors, proinflammatory cytokines, and environmental stress, activate the signaling cascade. MEK1 and MKK7 signal through parallel pathways, before converging to activate transcription factors (TF), such as Elk1, c-Jun and ATF2. These TF induce *COX-2* gene transcription. **(C)** qRT-PCR of *PP1A*, *HECTD4* and *COX-2* mRNA in control and *HECTD4*-KD cells, showing that *COX-2* mRNA levels rise upon *HECTD4* depletion. These cells have been used in the experiments shown in **Fig. S4D and Fig. 4B.** Error bars represent mean +/- SD. Significance was calculated using one-way ANOVA test, with Dunnett's multiple comparison test. **(D)** Trametinib was used to block MEK1 and investigate the effect of this inhibition on COX-2 activation. The qRT-PCR demonstrates that the drug fails to reverse *COX-2* mRNA induction in *HECTD4*-KD cells. Error bars represent mean +/- SD. Significance was calculated using unpaired student t-test. The baseline (no treatment) mRNA levels of *PP1A, HECTD4* and *COX-2* in cells used for this experiment are shown in **Fig. S4C.** The same cells were used in the experiments shown in **Fig. 4B. (E)** Western blot of the control and *HECTD4*-KD cells treated with trametinib. MEK1 is effectively inhibited, as demonstrated by the reduced activation of its direct target ERK 1/2 pT202/Y204. The increased COX-2 level in the *HECTD4*-KD is not affected by this inhibition, consistent with the qRT-PCR data shown in **Fig. S4D**. **(F)** qRT-PCR data demonstrates that MKK7 inhibition by the drug 5Z-7-oxozeanol (OXO) effectively reverses *COX-2* transactivation in *HECTD4*-KD cells. Error bars represent mean +/- SD. Significance was calculated using unpaired student t-test. **(G)** Western blot showing increased MKK7 protein in *HECTD4*-KD (sh #1 and sh #3) MDA-MB-231 cells growing in suspension. **(H)** *HECTD4* depletion does not impact *MKK7* mRNA levels, as shown by the qRT-PCR data from the same experiment in **Fig. S4A**. Error bars represent mean +/- SD. Significance was calculated using one-way ANOVA test, with Dunnett's multiple comparison test. (* $p<0.05$, ** $p<0.01$, *** $p<0.001$, **** $p<0.0001$)

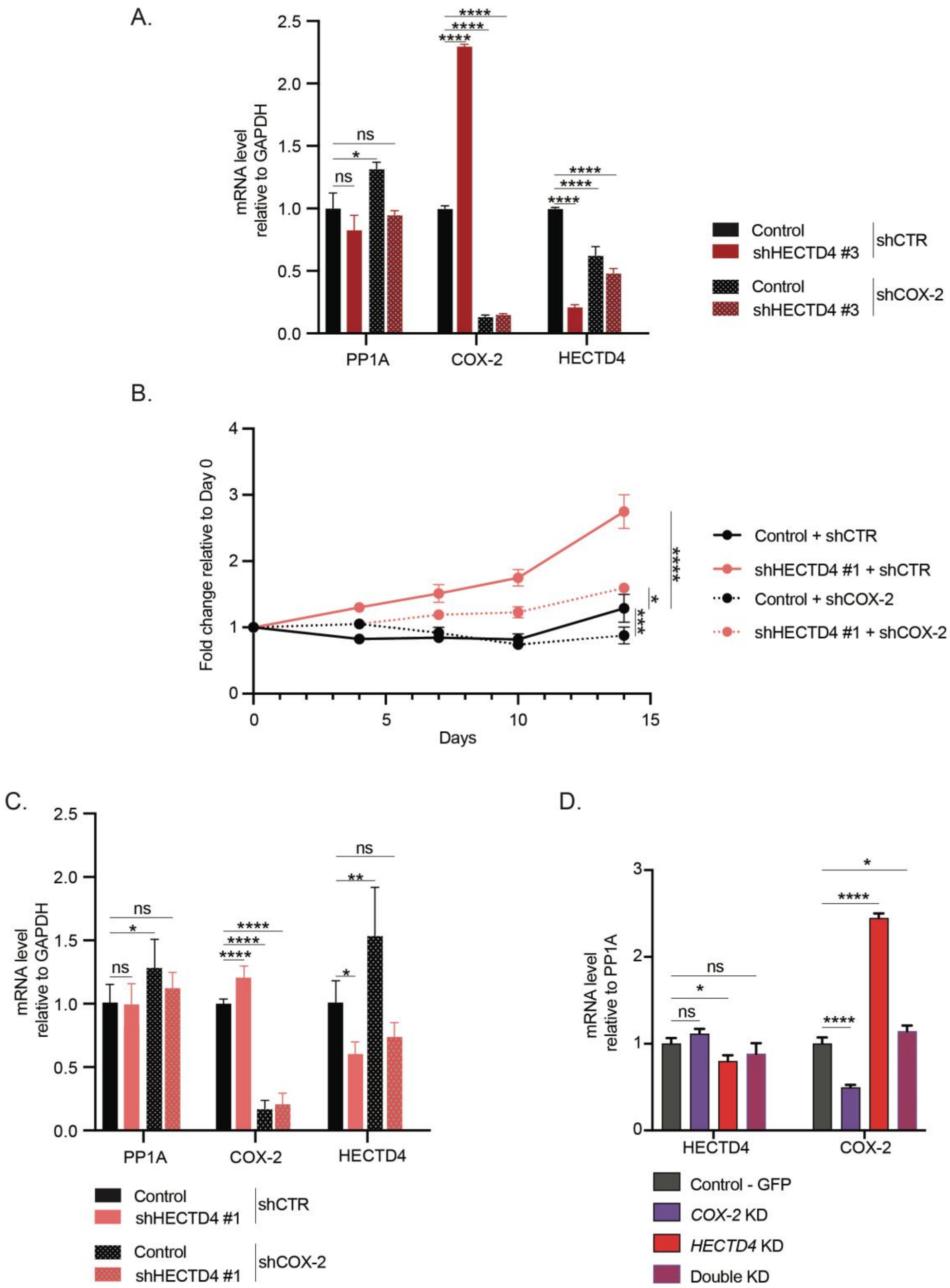
A.
mRNA level relative to GAPDH
2.5
2.0
1.5
1.0
0.5
0.0
ns
*
****
PP1A
COX-2
HECTD4
Control
shHECTD4 #3
shCTR
Control
shHECTD4 #3
shCOX-2
B.
Fold change relative to Day 0
4
3
2
1
0
0
5
10
15
Days
****
*
***
Control + shCTR
shHECTD4 #1 + shCTR
Control + shCOX-2
shHECTD4 #1 + shCOX-2
C.
mRNA level relative to GAPDH
2.5
2.0
1.5
1.0
0.5
0.0
ns
*
**
****
PP1A
COX-2
HECTD4
Control
shHECTD4 #1
shCTR
Control
shHECTD4 #1
shCOX-2
D.
mRNA level relative to PP1A
3
2
1
0
ns
*
****
HECTD4
COX-2
Control - GFP
COX-2 KD
HECTD4 KD
Double KD

**Fig. S5. HECTD4 modulation of COX-2-dependent proliferation *in vitro*** **(A)** qRT-PCR of *PP1A*, *HECTD4* and *COX-2* mRNA in control, *HECTD4*-KD, *COX-2*-KD and Double KD conditions. These cells were used in the experiments shown in **Fig. 4A.** Error bars represent mean +/- SD. Significance was calculated using one-way ANOVA test, with Dunnett's multiple comparison test. **(B)** Depletion of *HECTD4* compared to scrambled control increases cell proliferation under anchorage-independent, suspension conditions (ultra-low adherent culture dish). Depletion of *COX-2* in *HECTD4*-depleted cells reverts this phenotype. Cell viability and proliferation were measured by CellTiter Glo luminescence. Error bars represent mean +/- SD. Significance was calculated using with two-way ANOVA test, with Tukey's multiple comparison test (Day 14). Only cells harboring control and shHECTD4 #1 shRNA's are shown in this figure. Cells with control and shHECTD4 #3 shRNA's are shown in **Fig. 4A.** The mRNA levels of *PP1A, HECTD4* and *COX-2* in the cells used for this experiment are shown in **Fig. S5C. (C)** qRT-PCR of *PP1A*, *HECTD4* and *COX-2* mRNA in control, *HECTD4*-KD, *COX-2*-KD and Double KD conditions. These cells were used in the experiments shown in **Fig. S5B.** Error bars represent mean +/- SD. Significance was calculated using one-way ANOVA test, with Dunnett's multiple comparison test. **(D)** A mix of 2 shRNAs targeting *COX-2* efficiently reduced its mRNA level, both in the control and *HECTD4*-KD cells, reverting the HECTD4-mediated transactivation. These cells were used in the soft agar experiments shown in **Fig. 4C** and the *in vivo* mixing experiment shown in **Fig. 4D.** Error bars represent mean +/- SD. Significance was calculated using one-way ANOVA test, with Dunnett's multiple comparison test.

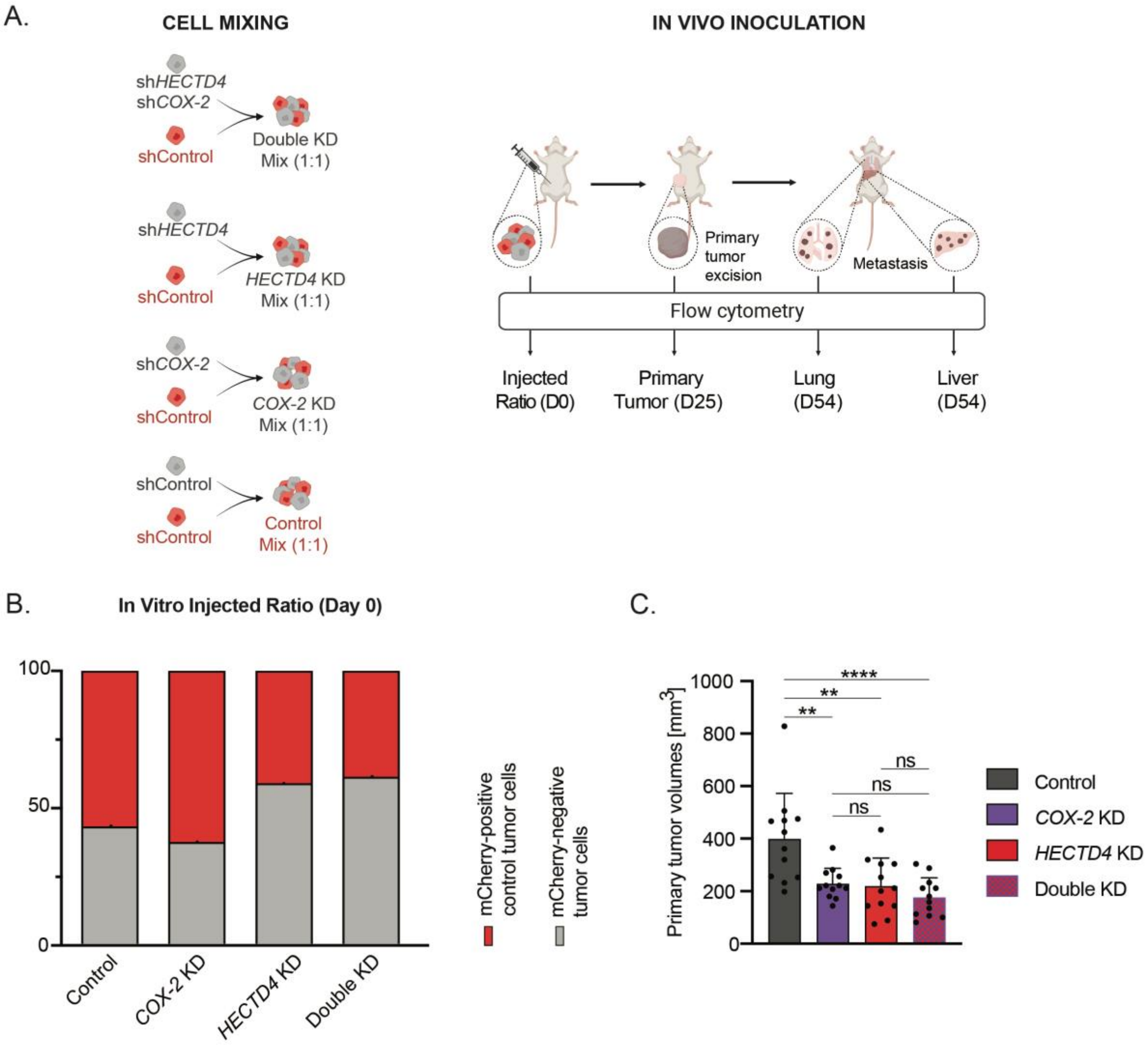
A.
CELL MIXING
shHECTD4
shCOX-2
shControl
Double KD
Mix (1:1)
shHECTD4
shControl
HECTD4 KD
Mix (1:1)
shCOX-2
shControl
COX-2 KD
Mix (1:1)
shControl
shControl
Control
Mix (1:1)
IN VIVO INOCULATION
Primary
tumor
excision
Metastasis
Flow cytometry
Injected
Ratio (D0)
Primary
Tumor (D25)
Lung
(D54)
Liver
(D54)
B.
In Vitro Injected Ratio (Day 0)
100
50
0
Control
COX-2 KD
HECTD4 KD
Double KD
mCherry-positive
control tumor cells
mCherry-negative
tumor cells
C.
Primary tumor volumes [mm3]
1000
800
600
400
200
0
****
**
**
ns
ns
ns
Control
COX-2 KD
HECTD4 KD
Double KD

**Fig. S6. HECTD4 modulation of COX-2-dependent tumorigenesis and metastasis *in vivo*** **(A)** Schematic representation of the tumor cells mixing experiment conducted *in vivo* to assess the effect of COX-2 depletion in *HECTD4*-KD cells. GFP+/ mCherry+ control cells (GFP-mCherry-shControl; *HECTD4*-WT and *COX-2*-WT, shown in red) and four GFP+/mCherry- conditions i.GFP-shControl (*HECTD4*-WT and *COX-2*-WT, shown in gray), ii.GFP-*COX-2*-KD (*HECTD4*-WT and *COX-2*-KD, shown in gray), iii.GFP-*HECTD4*-KD (*HECTD4*-KD and *COX-2*-WT, shown in gray), iv.GFP-Double-KD (*HECTD4*-KD and *COX-2*-KD, shown in gray) were combined in a ratio of 1:1 to give rise to four mixtures, respectively the control group, *COX-2*-KD group, *HECTD4*-KD group and double KD group. The mixtures were separately injected into the mammary fat pads of NSG mice. The mixtures were also seeded in vitro and analyzed by flow cytometry to ensure that each of the populations were equally represented in the mixture inoculated into the mammary fat pad at day 0. The primary tumors were resected at day 25 via survival surgery of the mice and the colonized organs – the lungs and livers – were harvested at day 54. The ratio of the different populations was analyzed by flow cytometry. **(B)** Fraction of the GFP+ /mCherry- tumor cells in the tumor cell mixtures at day 0 just before injecting into the mammary fat pads of NSG mice. The bars represent the percentage of red (GFP-mCherry-shControl) and green (i.GFP-shControl, ii.GFP-COX-2-KD, iii.GFP-HECTD4-KD, iv.GFP-Double-KD, shown in gray) cells. **(C)** The volumes of the primary tumors in the control, *COX-2*-KD, *HECTD4*-KD and double KD groups upon resection at day 25. Error bars represent mean +/- SD. Significance was calculated using with one-way ANOVA test, with Tukey's multiple comparison test. (* $p<0.05$, ** $p<0.01$, *** $p<0.001$, **** $p<0.0001$)

**Table S1.** List of target sequences of the short hairpins

| Gene | Manuscript name | Target sequence | Addgene # |
|---|---|---|---|
| HECTD4 | sh #1 | CGCTGCCTGTACCTTAGATTT | |
| HECTD4 | sh #2 | TGCGGAAGACACCCATATATA | |
| HECTD4 | sh #3 | GCGTCAGACACATTGACTATT | |
| COX-2 | sh #1 | GCTGAATTTAACACCCTCTAT | |
| COX-2 | sh #2 | CTATCACTTCAAACTGAAATT | |
| Control/Scramble | CTR | CCTAAGGTTAAGTCGCCCTCG | 136035 |

**Table S2.** List of guideRNA for ALFA introduction at the N-terminus of HECTD4

| Type | Sequence (including SpCas9 sgRNA scaffold) |
|---|---|
| pegRNA | 5'- GAAGCCTTCAGCCGGGGCCATGTTTTAGAGCTAGAAATAGCAAGTTAAAATAAG GCTAGTCCGTTATCAACTTGAAAAAGTGGCACCGAGTCGGTGCGCCGCCGCGG CCGCCGACGAGCCCTCGGTCAGTCTTCTTCTCAGCTCCTCCTCCAGTCTGCTCA TGGCCCCGGCTGTTTTTTT -3' |
| ngRNA | 5'- GCTCCAGTCTGCTCATGGCCCGTTTTAGAGCTAGAAATAGCAAGTTAAAATAAG GCTAGTCCGTTATCAACTTGAAAAAGTGGCACCGAGTCGGTGCTTTTTTT -3' |

**Table S3.** List of primers used for qRT-PCR.

| Gene | Forward primer | Reverse primer |
|---|---|---|
| GAPDH | 5'-CCTCAACGACCACTTTGTCAAG-3' | 5'-TGTTGCTGTAGCCAAATTCGTT- 3' |
| PP1A | 5'-CATCTGCACTGCCAAGACTGAG-3' | 5'-TCTTTCACTTTGCCAAACACCA- 3' |
| HECTD4 | 5'-AGGTACAGTGTGAACGGCTG-3' | 5'-AGAACATGCAGGCTCGAACA- 3' |
| COX-2 | 5'-CTGGCGCTCAGCCATACAG-3' | 5'-CGCACTTATACTGGTCAAATCCC-3' |
| MKK7 | 5'-GAAGGAGAAGCACGGTGTCA-3' | 5'-CTCCACCTGGGAGGGAGAC- 3' |

**Table S4.** List of antibodies used for Western Blot

| Protein/Target | Company | Cat# | Concentration |
|---|---|---|---|
| HECTD4 | ThermoFisher | PA5-61010 | 1:1000 |
| Ubiquitin | Cell Signaling | 14049S | 1:1000 |
| Vinculin | Sigma | MAB3574 | 1:1000 |
| ALFA tag | Nanotag Bio. | N1505-HRP | 1:3000 |
| β-actin | Cell Signaling | 3700S | 1:1000 |
| COX-2 | Cell Signaling | 12282S | 1:1000 |
| HA tag | Sant Cruz Bio. | SC-7392 | 1:200 |
| MKK7 | Cell Signaling | 4172S | 1:1000 |
| MAPK total (Erk1) | Cell Signaling | 4695T | 1:1000 |
| Phospho-MAPK (T202/Y204) | Cell Signaling | 4370T | 1:1000 |
| PCNA | Cell Signaling | 13110S | 1:1000 |
